# Pre and Post antibiotic epoch: insights into the historical spread of antimicrobial resistance

**DOI:** 10.1101/2024.09.03.610986

**Authors:** Adrian Cazares, Wendy Figueroa, Daniel Cazares, Leandro Lima, Jake D. Turnbull, Hannah McGregor, Jo Dicks, Sarah Alexander, Zamin Iqbal, Nicholas R. Thomson

**Author notes:** Corresponding author. (A.C.); (Z.I.); (N.R.T).

## Abstract

Plasmids are now the primary vectors of antimicrobial resistance, but our understanding of how human industrialisation of antibiotics influenced this is limited by a paucity of data predating the antibiotic era (PAE). By investigating plasmids from clinically relevant bacteria isolated between 1917 and 1954 and comparing them to modern plasmids, we captured over 100 years of evolution. We show that while all PAE plasmids were devoid of resistance genes and most never acquired them, a small minority evolved to drive the global spread of resistance to first-line and last-resort antibiotics in Gram-negative bacteria. They have evolved through complex microevolution and fusion events into a distinct group of highly recombinogenic, multi-replicon, self-transmissible plasmids that now pose the highest risk to resistance dissemination, and therefore human health.

**One Sentence Summary:** Pre-antibiotic era bacteria reveal the origin and evolution of drug-resistance vectors.

## Main Text

The emergence and spread of antibiotic-resistant bacteria through the use and overuse of antibiotics is one of the most significant side effects that human activity has imposed on our environment, perhaps rivalling climate change itself. As well as saving many millions of lives, the mass production and use of antibiotics has arguably been the most important selective pressure shaping bacterial evolution in the last 80 years. In many instances, selection has favoured highly transmissible multi-drug-resistant (MDR) bacteria. These MDR strains have disseminated across the world through globalisation: carried in human guts, through foods, animals and many other conduits. These now present an immediate threat to human health, with over 4.95 million deaths associated with antimicrobial-resistant infections in 2019 (*1*).

Originally, antibiotics were found to be naturally produced by bacteria and fungi as anticompetitive factors to kill competing microbes (*2*). In addition to the biosynthetic genes that produce antibiotics, bacteria can carry other genes that confer resistance to these antibiotics. In recent times, it is these antibiotic-resistance genes that have coalesced into discrete transferable units that have been widely shared amongst priority pathogens of humans and animals (*3*), explaining the current health crisis. Whilst we know a lot about antibiotic-resistant pathogens and the individual resistance genes, much less is known about the phylodynamics and evolutionary trajectories of the vectors upon which these antibiotic-resistance genes are transferred between bacterial hosts. Plasmids are one of the most important vectors of AMR genes, playing pivotal roles in the emergence of MDR pathogens and community and hospital outbreaks (*4, 5*). It is becoming increasingly common for a single bacterial cell to harbour single or multiple different plasmids that confer resistance to many, all frontline, or even last-line treatments (*3*). Despite their importance, the origins of resistance plasmids and how the anthropogenic introduction of antibiotics influenced plasmid evolution in the first place remain largely unknown. The answers to these questions have clear implications for understanding how antibiotic usage drives plasmid-mediated resistance; to address this, we need to understand the nature and composition of plasmids spanning the antibiotic epoch.

### Diversity of Murray plasmids and their relatives

To understand the life history of plasmids since the introduction of antimicrobials, we investigated the genomes (n=368) of a unique collection of pathogenic *Enterobacteriaceae* taken from patients between 1917 and 1954 (the “Murray Collection”) (*6, 7*), a key historical period spanning the discovery and first use of therapeutic antibiotics (Fig. 1A), to the current day.

**Fig. 1.**
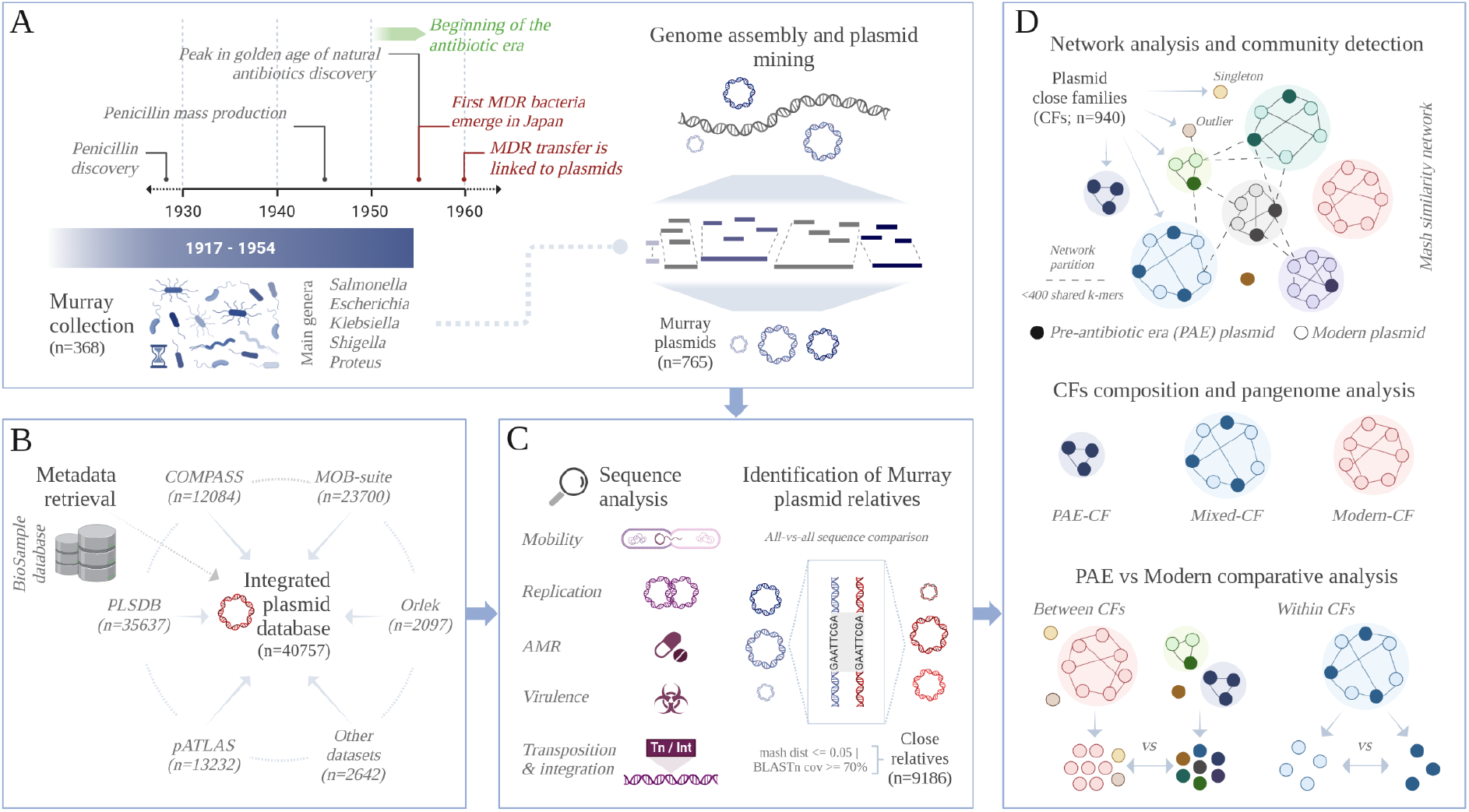
Framework for the analysis of PAE plasmids and their modern relatives. (**A**) Historical context and composition of the Murray collection, source of the PAE plasmids reconstructed in this study. (**B**) Collection of plasmid genomes integrated from multiple sources in the public archive and used for identifying Murray plasmid relatives. (**C**) Functional annotation of plasmid sequences and identification of Murray plasmid close relatives. (**D**) Schematic representation of plasmid close families (CFs) identified in the similarity network of Murray plasmids and their close relatives (top); their classification based on PAE and modern plasmids composition (middle); and the comparison between PAE and modern plasmids from the network (bottom). PAE (<=1954) and modern (>1954) plasmids are depicted as filled or open circles across the figures in panel D, respectively. Plasmids from Modern-CFs were compared against all PAE plasmids in the network (bottom-left), whereas modern plasmids from Mixed-CFs were compared against PAE members of the same close family (bottom-right).

We generated sequences for 765 “Murray Collection” plasmids. They were identified in isolates from 6 genera and 9 species (fig. S1 and table S1). We compared these sequences to 40,757 plasmid genomes retrieved from all available plasmid databases in the public archive, showing that 9,186 plasmids were significantly similar (defined as 70% BLAST coverage of the Murray plasmid or being a mash distance of at most 0.05 apart) to 715 Murray Collection plasmids (Fig. 1A-C).

The 9,951 Murray and related plasmids range in size from ∼0.5 to 606 kb, are globally distributed across at least 79 countries from 6 continents, and are linked to more than 72 species and 25 genera (fig. S2 and table S1). Notwithstanding this, most (95%) of them were identified in 5 genera: *Escherichia, Shigella, Klebsiella, Salmonella* and *Enterobacter*, all deemed global priority pathogens by the WHO due to their high resistance to last-line antibiotics (*8*).

We constructed a sequence-similarity network to determine the diversity and relationships between this 9,951-plasmid collection and performed a community detection analysis to define groups of evolutionarily related plasmids. Here, relatedness is determined by a mash similarity threshold of >=400 shared k-mers used to partition the network (Fig. 1D and fig. S2). This resulted in the detection of 940 network communities, hereafter referred to as close families (CFs) (table S2). These data showed a strong population structure: 22 CFs encompass 50% of the plasmids in the network. On the other hand, 264 CFs are disconnected from all others and represent singletons (Fig. 2A).

**Fig. 2.**
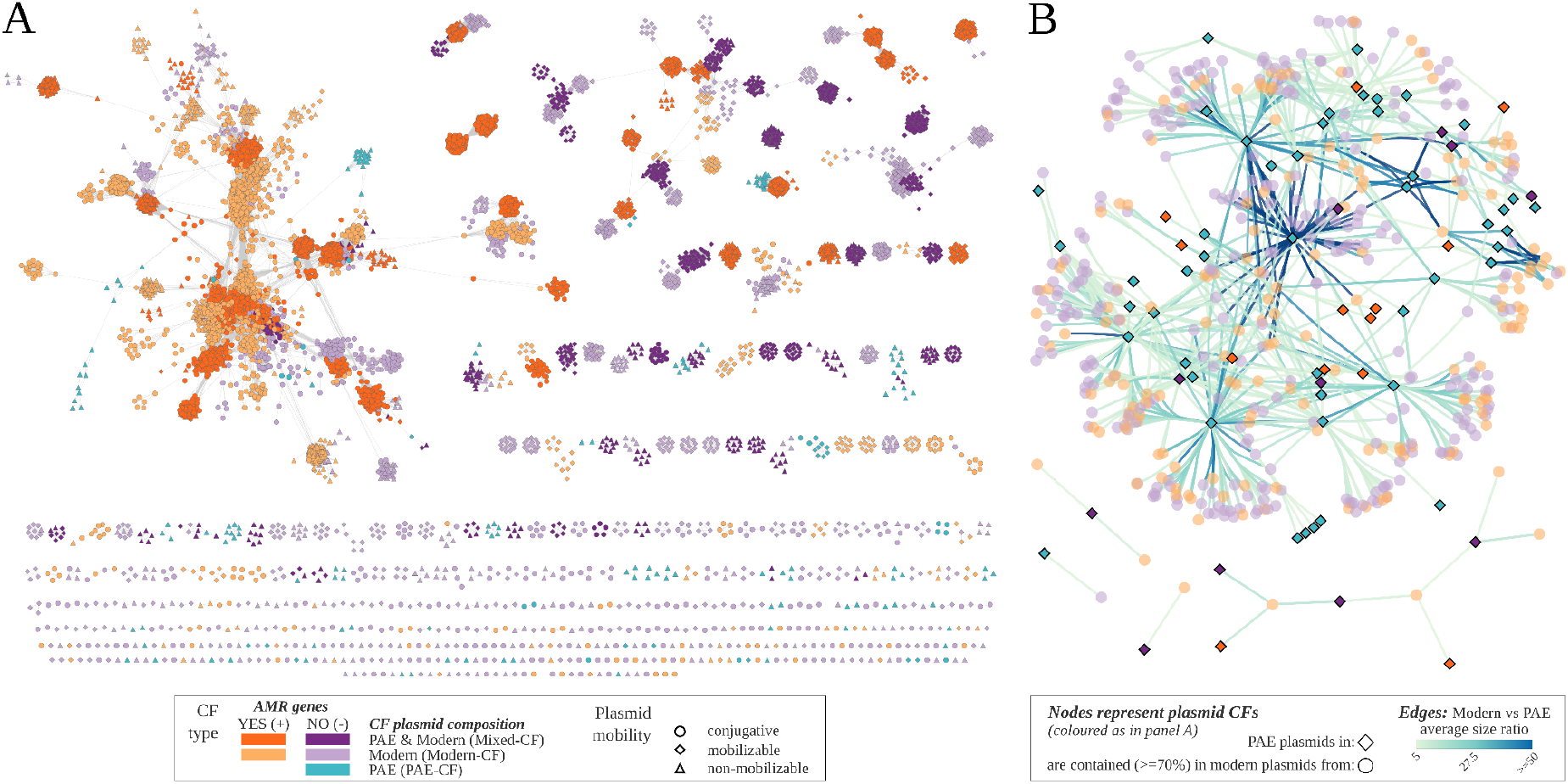
Diversity of PAE and related modern plasmids and their contribution to AMR. (**A**) Plasmid similarity network. Plasmids are represented as nodes in the network, connected by edges if their genomes share a minimum mash distance of 0.05. The plasmids are coloured based on the type of CF they were classified into, denoting the composition in terms of PAE and/or modern plasmids and the presence of resistance plasmids. The shape of the node is determined by the plasmid’s mobility classification. (**B**) PAE plasmid containment network. The nodes in the network represent plasmid CFs, collapsed from the similarity network shown in panel A and coloured accordingly. The node’s shape differentiates Modern-CFs (circles) from other CF types (diamonds). Nodes are connected by an edge if at least one plasmid from a Modern-CF was identified as containing (=>70 coverage) one or more Murray plasmids via BLAST. The edges’ colour reflects the average size difference between the modern and contained PAE plasmids. To reduce complexity, the network illustrates interactions where the modern plasmid is, on average, at least 5x larger than the contained PAE plasmid (59% of Modern-CF plasmids); all interactions are described in table S4.

### Composition and evolution of plasmid close families

To understand how well plasmids collected before the widespread use of antibiotics were represented across the network, those isolated on or before 1954 (the last year sampled in the Murray collection) were denoted ‘pre-antibiotic era’ plasmids (PAE, comprising 22 public archive sequences plus 765 Murray plasmids). All remaining plasmids were labelled modern-era plasmids or Modern (dates 1956 - 2021, 99% collected after 1990; see Methods; fig. S2 and table S1).

From Fig. 2, it is clear that within the network, there are three types of CF based on the sampling dates of the plasmids (see Fig. 1D for a schematic representation): those exclusively represented by PAE (PAE-CF; n=103), or Modern plasmids (Modern-CF; n=749) and the third comprising a mixture of both PAE and Modern plasmids (Mixed-CF; n=88). Our pangenome analysis of the plasmid CFs identified a core genome in 97% of them (excluding those comprising one plasmid; fig S3 and table S3), thus indicating that plasmids in Mixed-CFs have maintained a recognisable clonal frame over considerable time frames — the longest in our dataset being between 1917 and 2020, and over the period spanning the introduction and widespread use of antibiotics. What is more, plasmids within this CF category are the most common across our network (Fig. 2A and S4), representing 60% of all plasmids studied here and 77% of all PAE plasmids. The remaining 23% of PAE plasmids, which comprise the PAE-CFs (n=184), have not been seen again in any publicly available plasmid database since 1954, using these search criteria.

Next, we investigated the plasmids in Modern-CFs. Although Mash is highly efficient for clustering large-scale genomic datasets, the algorithm is sensitive to high sequence divergence and large size differences between the sequences being compared (*9*). Given all of the 9,186 plasmids from databases in our network had shown some similarity to the Murray collection plasmids by BLAST, we repeated this approach to define the Modern-CF plasmids relationship with PAE plasmid sequences (Fig. 1D). Our analysis revealed that 98% (n=3,646) of the Modern-CF plasmids contained sequences matching at least one of 506 PAE plasmids (defined as covering at least 70% of a reference PAE plasmid by BLAST; see methods, table S4). The principal explanation for the low Mash scores was a large difference in size between the Modern and PAE reference plasmid. Only 6.6% (n=241) of Modern-CF plasmids displayed similar size to their matching PAE plasmid (+/- 30% size difference) and were not linked to it by Mash because they showed high sequence divergence. The other 93.4% of Modern-CF plasmids are, on average, 12 times larger than the PAE reference plasmid they contain, with 59% of these being at least 5x larger (table S4 and fig. S5).

The identification of numerous PAE plasmids subsumed into larger Modern-CF plasmids suggests that the latter were predominantly formed through the fusion of multiple unrelated sequences. Consistent with this, 58% of the Modern-CF plasmids carry multiple plasmid replication genes, *rep* (fig. S6 and table S1). These genes are used for categorising plasmids into incompatibility (Inc) groups (or “Inc types”), with one Inc type traditionally representing a plasmid type (*10*) – thus, multi-replicon plasmids are strongly indicative of having previously been through a fusion event. What is more, a network illustrating PAE plasmid containment between CFs showed that a given PAE plasmid sequence can be contained in plasmids from multiple different Modern-CFs and, in turn, plasmids in these Modern-CFs can contain multiple different PAE plasmids (Fig. 2B and fig. S7). This complex pattern of recombination results in 69% of the Modern-CFs having plasmids with unique gene and sequence profiles (defined as CFs comprising a single plasmid; fig. S4), with 31% disconnected from all other CFs.

### The relevance of AMR to plasmid evolution over the last decades

To understand the relative contribution of PAE plasmids and their Modern relatives to the dissemination of AMR, we searched for all known AMR genes. We identified 21,378 instances of more than 300 different AMR genes encoded in 38% (n=3,815) of the plasmids in the network (table S5 and fig. S8). No AMR genes were detected in PAE plasmids. By contrast, the number of resistance genes across their modern relatives ranged from 1 to 40, many of which confer resistance to broad-spectrum and/or last-resort antibiotics, including carbapenems, tigecycline and polymyxins such as colistin. Of note, the catalogue of resistance genes identified includes most of the known extended-spectrum beta-lactamase gene families (table S5 (*11*)). The most striking combinations were resistance to 13 drug classes in the plasmid pHZ003 (NZ_MN476094.1) and 8 classes in the plasmid pSY3626C1 (NZ_CP059044.1). In terms of the number of AMR genes, the most extreme plasmids encoded either 40 or 38 individual resistance genes linked to gene or AMR cassette duplications (fig. S7).

Next, by mapping the occurrence of AMR genes across the network, we determined the relative contribution of different CFs towards the acquisition and dissemination of drug resistance (Fig. 2A, 3B and S8). Only 25% of CFs included plasmids with known AMR genes (table S2 and fig. S4), with the average number of resistance genes varying considerably amongst CFs, from 0.005 to 13 (excluding CFs with a single plasmid), whereas the frequency of resistance plasmids within a CF ranged from 0.5 to 100%. Evident from the network (Fig. 2A) is that AMR carriage is independent of the CF’s plasmid composition: 39 Mixed-CFs and 203 Modern-CFs contain resistance plasmids. Nevertheless, despite consisting of only 22% of the total plasmids in the network, plasmids in Modern-CFs carry 50% of all the AMR genes detected and, on average, more AMR genes than those in Mixed-CFs (Fig. 3B and fig. S8).

**Fig. 3.**
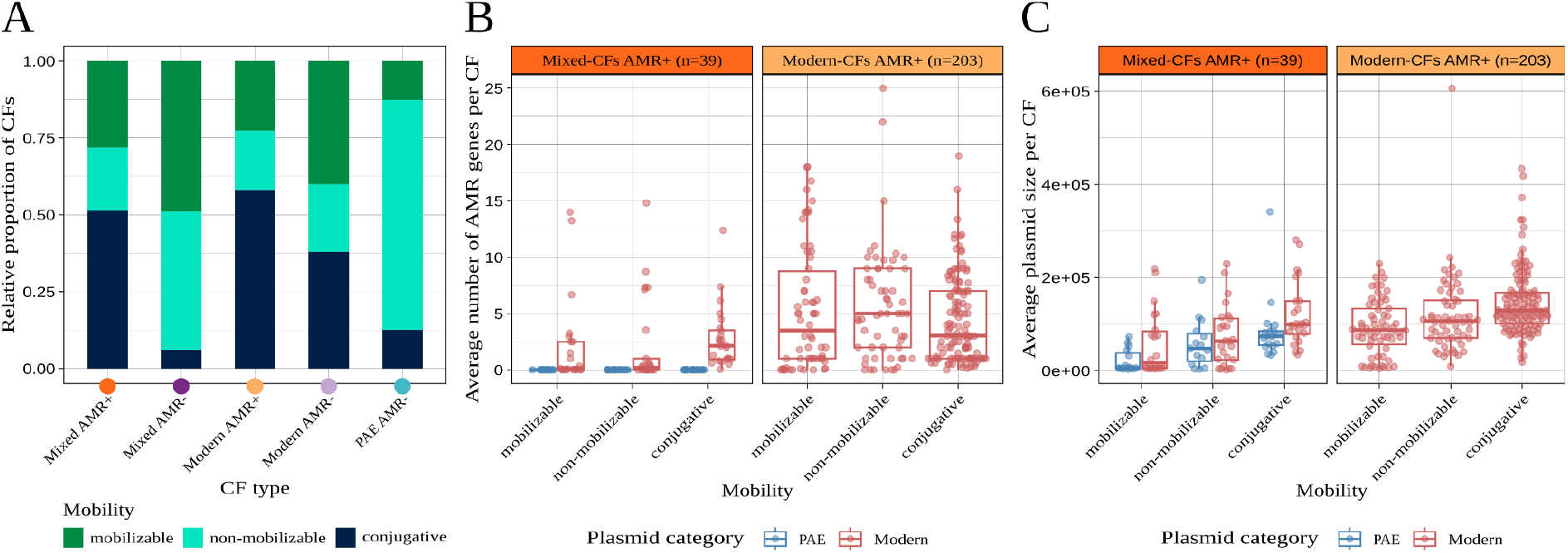
Mobility, AMR gene carriage and size distribution across plasmid CFs. (**A**) CFs plasmid mobility classification. The plot is arranged by CF type (X-axis; a coloured dot matches the categories described in Fig. 2A) and shows the proportion of CFs within a given CF type classified as conjugative, mobilizable or non-mobilizable. The classification was determined by the most frequent mobility category identified in the CF (see fig. S9). (**B**) AMR gene content across plasmid CFs. The plots are split by CF type, and within each plot, values are divided by mobility type and PAE or modern plasmid for Mixed-CFs (left). Each point in the plots represents a CFs’ average value. (**C**) Average plasmid size across CFs with resistance plasmids. Plots and data points are structured as in panel B. The distribution of average plasmid size for all CFs is shown in fig. S2.

### The link between plasmid size, their pangenome, mobility traits and AMR

To understand if there was a link between AMR and mobility, plasmids were divided into three mobility classes: conjugative (self-transmissible), mobilizable (needing components from helper plasmids), and non-mobilizable (table S1 and fig. S8 (*12*)). Our data showed that 58% and 51% of the Modern-CFs and Mixed-CFs containing resistance plasmids were dominated by conjugative plasmids, respectively (Fig. 3A and table S2). By contrast, this fell to 38% and 6% among Modern-CFs and Mixed-CFs with no resistance genes, respectively. Here, combined mobilizable and non-mobilizable represented from 62 to 94% of the plasmid mobility types. Across all CFs, conjugative plasmids exhibit the largest average size (Fig. 3C and fig. S2), and they carry the most AMR genes among Mixed-CFs (Fig. 3B). When splitting out by CF types, it is also clear that Modern-CF plasmids are on average larger and carry more AMR genes compared to the same mobility type in Mixed-CF plasmids (Fig. 3B and C). Furthermore, our data showed that 61% of all conjugative plasmids (n=5,430; 48% of conjugative in Mixed-CF and 84% in Modern-CF; fig. S6) are multi-replicon and that they constitute 78% of all multi-replicon (n=4,256) plasmids identified in the network (table S1). The mobilizable and non-mobilizable types only represent 8% and 14% of multi-replicon plasmids, respectively (fig. S6).

Uniquely, by comparing the PAE plasmids to their conserved modern descendants in the 88 Mixed-CFs, we were also able to look at the evolutionary trajectory of plasmids that have maintained a clonal frame during the antibiotic era. In addition to plasmid size, the presence or absence of AMR genes and plasmid mobility functions, we characterised the plasmids’ pangenome composition and searched for the presence of genes involved in virulence (table S6) and selfish mobile genetic elements linked to AMR dissemination, namely integrons and transposons (table S1).

Comparison at the CF level (Fig. 4A) revealed that the average size of plasmids in Mixed-CFs containing at least one resistance plasmid (AMR+) was markedly larger than in Mixed-CFs with no resistance plasmids identified (AMR-; 71.4 kb vs 21.7 kb; *P* = 1.292e-09). This observation was strongly correlated with AMR gene content, the presence of transposases (mean 5.9 vs 2.5; *P* = 7.475e-11) and/or integrons (none detected in AMR-) and, consistent with the above, the mobility type: plasmids encoding conjugative machinery (typically comprising ∼12 to 32 genes (*12*)) are the prevalent type in more than half of AMR^+^ CFs. Conversely, AMR^-^ CF plasmids were predominantly mobilizable (49% of CFs) or carried no detected mobility functions (45% of CFs; Fig. 4A). Our pangenome analysis also showed that AMR+ CFs tend to feature a smaller core genome compared to their AMR-counterparts, expressed both as the proportion of core genes in the pangenome (Fig. 4A; mean 12.3% vs 39.3%; *P* = 1.292e-09) and the average number of core genes per plasmid (fig. S3). This difference, indicative of greater genome flexibility and cargo capacity in AMR+ CFs, was also evident among Modern-CFs (fig. S3). There was also a significant difference in virulence genes content between AMR+ and AMR-Mixed-CF groups, albeit smaller than that determined for the other features (mean 1.6 vs 1.2; *P* = 1.972e-10).

**Fig. 4.**
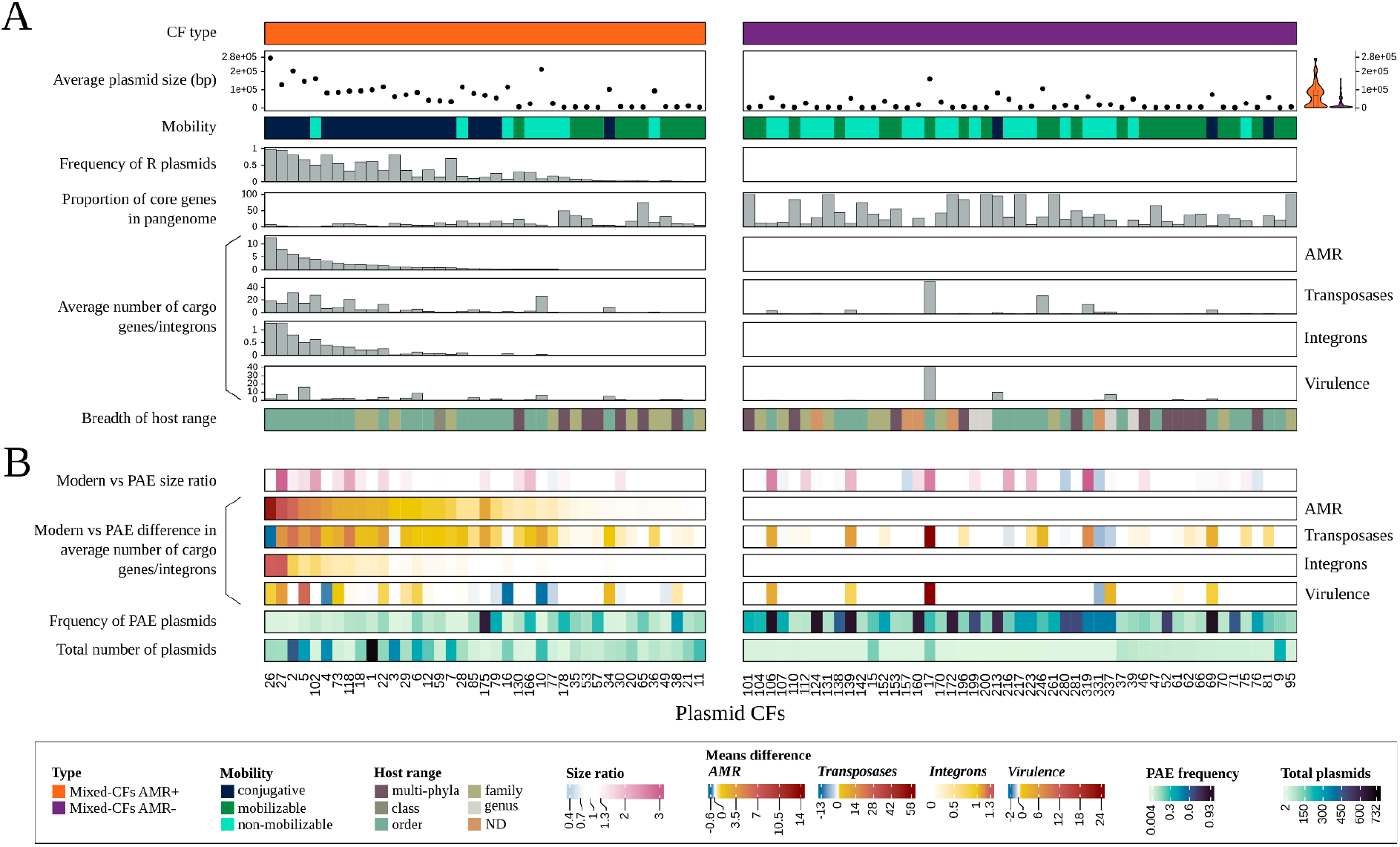
Evolution of PAE plasmids that have maintained a clonal frame. (**A**) Comparison of plasmid characteristics across Mixed-CFs. The 88 Mixed-CFs identified in the plasmid similarity network are indicated over the X-axis, arranged from higher to lower average number of AMR genes in the CF (left to right). The plots are split by CF type, reflecting the presence (left, orange) or absence (right, purple) of resistance plasmids in the CF. Mobility and host range were defined by the most frequent category identified in the CF. (**B**) Differences between modern and PAE plasmids within Mixed-CFs. The four top rows in the panel summarise differences in size and number of cargo genes expressed as the ratio of average values, using modern plasmid values as the numerator. The significance of modern vs. PAE differences was assessed by implementing a sampling with replacement strategy, illustrated in fig. S10 (results are detailed in table S8).

The comparison of PAE and Modern plasmids within a Mixed-CF showed that AMR genes are carried by the latter, as previously indicated. What was surprising, however, was that plasmid evolution in Mixed-CFs was not primarily via gene accumulation and accretion, which would have resulted in more recent plasmids being larger than their PAE counterparts. Fig. 4B shows that in most (67%; n=59) Mixed-CFs, PAE and Modern plasmids display a comparable average size (+/- 30% size difference); we refer to this as size stasis. For the remaining Mixed-CFs, Modern plasmid’s average size was larger (up to 3x; 28% of CFs; n=25) or smaller (4% of CFs, n=4) than their PAE counterparts (Fig. 4B and fig. S10). Despite the large proportion of Mixed-CFs identified in size stasis, our analysis revealed that various Modern plasmids have acquired Integrons, Transposase, AMR or virulence genes when compared to their reference PAE backbones across different CFs (Fig. 4B). Notably, we also found that, unlike AMR genes, those encoding virulence factors or transposases were common in PAE plasmids. In line with this finding, we detected a few cases of CFs where the average number of virulence or transposase genes was lower in Modern plasmids. Overall, the acquisition of AMR, transposase genes and Integrons correlated well among AMR+ CFs (Fig. 4). The patterns of size and cargo gene difference between PAE and Modern plasmids across Mixed-CFs indicate that gene gain, loss, and replacement have all shaped the evolution of PAE plasmids that maintained a clonal frame during the antibiotic era.

### Beyond Murray plasmids

Finally, our initial search for Murray plasmid relatives (Fig. 1C) revealed they were significantly similar (>= 70% coverage) to 22% of the plasmids in public databases (n=40,757); we consider these to be Murray close relatives. Given our observations from the Modern-CFs above, we relaxed our overall similarity threshold to as low as 1% plasmid coverage. In doing so, we were able to link the Murray plasmids to 38% (n=15,644) of all reported plasmids in the public archive (fig. S11), uncovering a subset of Murray extended relatives (n=6,626) which contain fragments of Murray plasmid sequences below the initial coverage threshold used to identify close relatives. On the other side of the spectrum, we determined that 62% (n=25,113) of the plasmids in the public archive have no similarity to any Murray plasmid (defined as displaying 0% coverage of a Murray plasmid sequence; table S7; Fig 5 and fig. S11). We refer to these plasmids as Murray-unrelated.

**Fig. 5.**
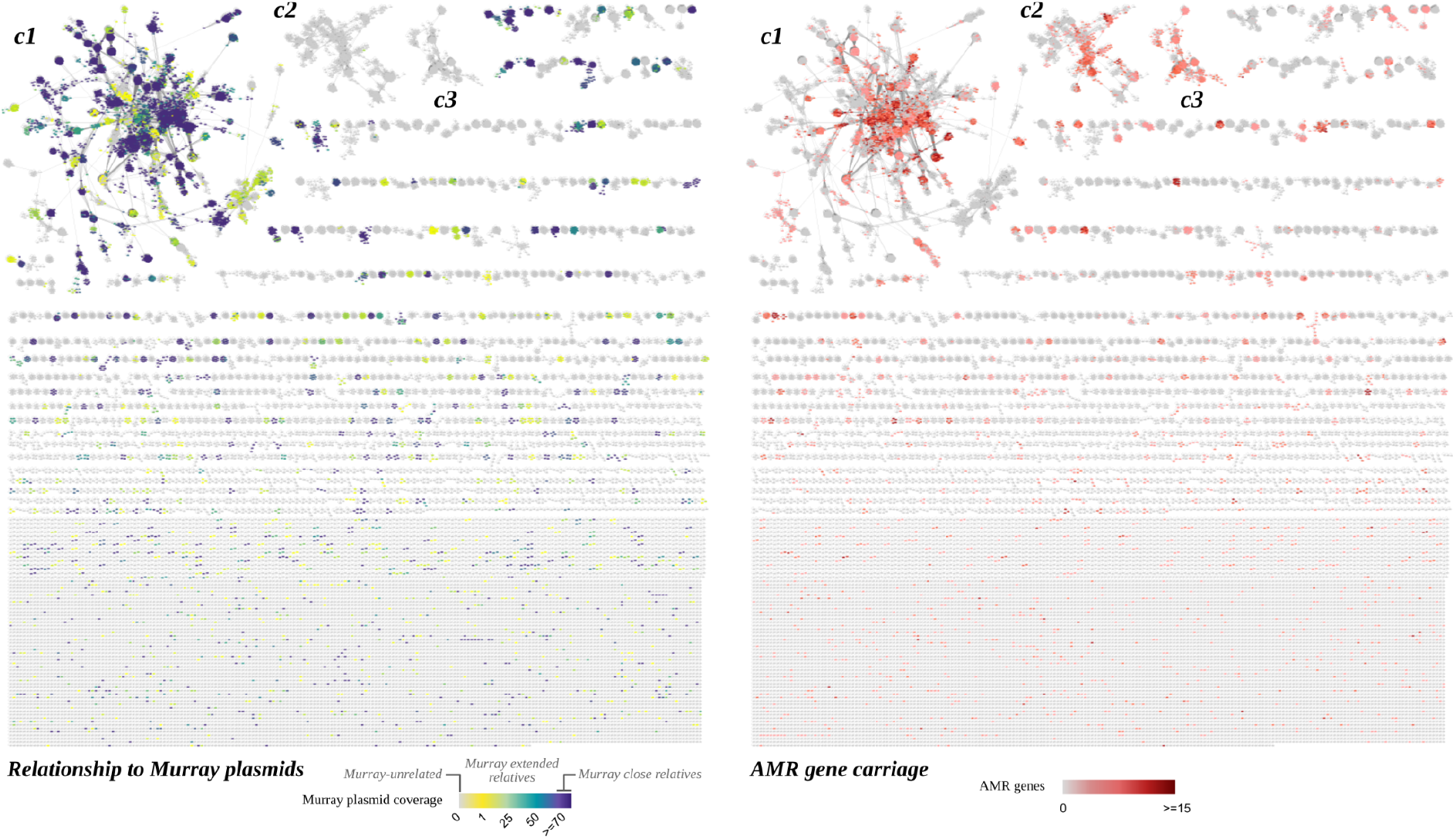
Landscape of known plasmids and their links to Murray plasmids and AMR. Similarity network of 40,757 plasmid genomes collected from the public archive (see Fig. 1B). Plasmids are represented as nodes in the network, connected by edges if their genomes share a minimum mash distance of 0.05. The node colour denotes similarity to Murra plasmids (left; expressed at the coverage of the best match identified via BLAST) or AMR gene content (right). The three largest connected-component clusters are labelled *c1* (n=12556; 95.3% *Enterobacteriaceae*), *c2*(n=1222; 99.6% *Enterococcaceae*) and *c3*(n=707; 99.7% *Staphylococcaceae*).

To reveal the range of plasmid diversity described by Murray close and extended relatives, we created a full similarity network combining the 40,757 plasmids from our integrated database (Fig. 5). Evident from this network was a single dominating cluster notable for the Murray plasmid fragments and AMR genes it contained (labelled *c1* on Fig. 5). In fact, about 36% of the 58,870 AMR genes we detected across this network are carried by Murray close relatives (table S7). Of note, if we rank all plasmids in terms of the number of AMR genes they carry, then from the top 100 with the most AMR genes, 92% are contained in the largest dominating network cluster, and 75% are related to a Murray plasmid (38% are Murray close relatives).

Murray close relatives and Murray-unrelated plasmids show a strong difference in host taxonomy (fig. S11). The former group comprised two known taxonomic classes, *Gammaproteobacteria* and *Actinomycetia*, with the former representing 98% of the sequences. This fraction decreased to 33% among Murray-unrelated plasmids, with Bacilli accounting for 29% as the second predominant class. Overall, the hosts of the Murray-unrelated plasmids comprised 48 taxonomic classes.

Despite the differences in host taxonomy and the lack of detectable relationship to Murray plasmids at the DNA level, we found that Murray-unrelated plasmids display common characteristics in terms of population structure and AMR gene distribution (Fig. 5), i.e. tight groups of closely related plasmids are clearly distinguishable (sometimes embedded inside large complex clusters), and AMR genes concentrate in a minority of discrete plasmid groups. These features are best exemplified by the two second-largest clusters in the network (Fig. 5), which exhibit a high incidence of AMR genes and are formed almost exclusively by plasmids linked to the families *Enterococcaceae* (n=1,222; *c2*) and *Staphylococcaceae* (n=709; *c3*). Among these plasmids, only one displays similarity to a Murray plasmid sequence (28% coverage; table S7).

We then explored whether evolutionary patterns linked to AMR carriage among Murray close relatives were shared with Murray-unrelated plasmids. We observed a higher proportion of resistance versus non-resistance plasmids among conjugative and multi-replicon plasmids and a higher average number of AMR genes in multi-replicon backbones of Murray-unrelated plasmids. This is in line with our findings on Murray close relatives (fig. S12); however, the link between AMR carriage and conjugative plasmids was not as strong, and here ‘conjugative’ was not the prevalent mobility platform for multi-replicon plasmids. These differences are clearer when focusing on the large *Enterococcaceae* and *Staphylococcaceae* network clusters, compared to the unrelated Gammaproteobacteria plasmids (fig. S12). The difference is likely explained by the known lower incidence of conjugative plasmids reported in Gram-positive bacteria compared to Gram-negative taxa (*13*).

## Discussion

Plasmids are known to be primary vectors of AMR genes globally (*4, 5*). Our work was inspired by E.G.D. Murray and the unique collection he put together, as well as the seminal work on the experimental characterisation of the Murray collection by Naomi Datta and Vicky Hughes that was published in 1983 (*6, 14*). Through access to this unique collection of historically relevant bacteria and genomics, this study unlocks past plasmid diversity to understand the origin and evolution of resistance vectors in the modern era. We show that key plasmid backbones, which have likely been residing in human pathogens for millennia, have played critical roles in the evolution and spread of multidrug resistance. The lack of known AMR genes in these historical plasmids provides the first molecular estimates for the emergence of resistance vectors to a short period during the antibiotic era. The plethora of resistance genes we identified in their modern relatives showcases how these pre-existing plasmid backbones have been pivotal for bacterial survival throughout the antibiotic epoch.

Our network analysis of the PAE plasmids reported in this study and their modern relatives identifies three classes of network community (Close Family, CF). PAE-CFs are formed exclusively by PAE plasmids that do not have descendants in public plasmid archives, likely because they are in undersampled or rare species, have been overlooked in current genomes, or possibly because they are extinct. Mixed-CFs encompass 60% of the plasmids in our network and comprise plasmid lineages that have maintained a recognisable clonal frame since the pre-antibiotic era, shaped by microevolutionary processes such as mutation, gene loss, gain and replacement. Modern-CFs constitute the remaining 37% of plasmids in the network and correspond to modern plasmids linked to PAE plasmid ancestors primarily via the absorption of one or more of their backbones; hence, they have arisen through fusion events resulting in conspicuously large plasmids. We show that resistance plasmids emerged in both Mixed-CFs and Modern-CFs and discuss shared characteristics below.

We found that fusion, which we use in a broad sense to encompass different possible molecular mechanisms, as an evolutionary process for plasmids is not a minor character but, in fact, should be centre-stage. It has long been known that plasmids were able to form hybrids (*15*); however, this was not considered a common occurrence or an important process on an evolutionary scale, but rather something that could happen under strong antibiotic selection in the lab. Our identification that most (66%) of our PAE plasmids were at some point subsumed into diverse chimeric plasmids, inverts this perception. We do not have the data to estimate how frequently fusion events happen, but clearly over a 100 year timescale, the descendants of such events have frequently been able to survive, which is in line with the ∼20% multi-replicon incidence observed in some plasmid datasets (*16*). This suggests that, at least over these timescales, we should model plasmid evolution quite differently from how we model normal species, as the child of a fusion event is a very different “DNA organism” to either parent - potentially much larger, and under different selective pressures due to differing genetic cargo. Finally, since 69% of Modern-CFs correspond to plasmids that do not cluster with other sequences, our data implies that a large proportion of these plasmid fusions are unique instances from a vast number of possibilities, through which the resulting plasmids have lost their clonal frame. The commonplace existence of this phenomenon is remarkable – equivalent to the frequent spontaneous birth of fertile hybrids between different species with different traits, as if lions and eagles frequently gave birth to fertile griffins.

Plasmid fusion, typically referred to as cointegration, is not exclusive to the antibiotic era or AMR carriage; e.g. several PAE plasmids feature multiple replication genes (indicative of prior fusions) and plasmids in 546 Modern-CFs do not carry known AMR genes. Nevertheless, our data shows that fusion/multi-replicon plasmids have played a dominant but overlooked role in the spread of resistance over the last decades: resistance plasmids in Modern-CFs account for half of the AMR genes identified in the network despite representing around one-fifth of the total plasmid sequences. This is likely due to their versatility and ability to subsume multiple unrelated plasmids into a single entity collecting and disseminating MDR, among other traits. In our collection of PAE and related modern plasmids, a self-transmissible lifestyle is the preferred evolutionary platform to build composite plasmids.

Our identification of resistance genes in 242 plasmid CFs of different composition types highlights that resistance vectors have emerged via diverse genetic backbones and mechanisms. Plasmids are not constrained to a single evolutionary path since the same PAE plasmid backbone can follow contrasting evolutionary strategies, namely microevolution or plasmid fusion. Notably, however, we found that resistance plasmids from Mixed-CFs and Modern-CFs share characteristics; for example, high genome plasticity and multi-replicon prevalence. Considering multi-replicon plasmids from both CF types originating through plasmid fusion, their difference might be explained by size constraints (i.e. the unrestricted growth in average size we see in Modern-CF plasmids is only permissible in certain backgrounds). Alternatively, Modern-CF plasmids are only a snapshot sampled in a relatively small time window, and may eventually collapse back to being recognisable Mixed-CF plasmids.

Whilst humans did not invent antibiotics, our study shows how the industrialisation and widespread use of antimicrobials as therapeutic agents changed the role and genetic identity of diverse plasmid lineages along with the way bacteria evolve resistance. We have shown that despite the high diversity of plasmid backbones seen in the Murray Collection, only a small subset of them (26% of the CFs linked to Murray plasmids) have acted as vectors for antimicrobial resistance. We also show that there are consistent themes underlying the different trajectories leading to AMR carriage. Along with the large cargo capacity and concomitant accumulation of intracellular mobile genetic elements, especially transposons, the enrichment of conjugative plasmids and those featuring multiple replication genes is prevalent among resistance vectors, particularly in *Enterobacteriaceae*. These characteristics can serve as a genetic barcode that identifies plasmids posing a higher risk for the spread of AMR. We identified specific highly recombinogenic and self-transmissible plasmid lineages that have been evolutionary successful as MDR vectors and should be monitored. Circumstances where multiple rearranged fusions of AMR plasmids are being generated are of particular concern to public health.

## Materials and Methods

### Murray plasmids

Three-hundred seventy genomes of strains from the Murray collection were assembled from Illumina reads (accession numbers are indicated in table S1; (*7*)) using unicycler v0.4.7 (*17*) with default options. Assembly metrics were calculated using quast v4.5 (*18*). Two assemblies resulting in <8 kb (M666 and M639) were removed from further analyses. Taxonomic classification of the remaining 368 genomes was performed with kraken2 v2.1.2 (*19*) and bracken v2.5 (*20*) on both the sequencing reads and assembly contigs. Kraken2 was run using the standard 8GB database (2021) and with the options “--use-names” and “--report” to output the taxa scientific names along with the taxonomy ID and a sample report, respectively. Bracken was run with default options. Taxonomy labels were consistent between the output from reads and contigs.

Depletion of chromosomal contigs from the assemblies was carried out with the script plasmidverify.py (*21*), which uses hmmsearch from HMMER 3.3 (*22*) and the database Pfam-A v33.0 (http://ftp.ebi.ac.uk/pub/databases/Pfam/releases/) to detect contigs of plasmidic origin based on gene content. The script was run with default settings. Contigs not circularised by unicycler but classified as plasmidic by plasmidverify were processed for circularisation with circlator v1.5.5 (*23*). The following settings were adjusted to run the circlator’s command “all” on the input Illumina reads: “--b2r_min_read_length 50 --merge_min_length 100 --merge_min_length_merge 200 --assemble_spades_k 107,97,87,77,67,57,51 --b2r_discard_unmapped --clean_min_contig_length 100”. The resulting sequences were combined with those circularised by unicycler and subsequently typed with plasmidfinder (*10*) (via abricate v1.0.1 (*24*); identity (--minid) and coverage (--mincov) cut-offs set to 80 and 60%, respectively) and mob_typer (from the mob_suite v3.0.1 (*25*); ran with default options). The sequences were further processed for plasmid identification by combining two approaches:

#### Mapping

The sequences were aligned against plasmids from the core subset of the integrated dataset generated for this study (see below) using minimap2 v2.17-r941 (*26*). We used the preset flag (-x) to tune the parameters for “assembly to reference alignment (asm5)”. Coverage of the query and reference sequences was calculated from the alignment with pafpy v0.1.1 (*27*). The query sequences were classified as plasmid or putative plasmid based on their similarity to known plasmids (coverage of the reference plasmid sequence and size ratio), circularisation, and presence of regions related to plasmid replication and/or mobility. The criteria used for classification are illustrated in fig. S13. Putative plasmid sequences were further examined by comparing them against the sequences available in the GenBank nucleotide collection (nr/nt) in June 2021 using BLAST online with default settings (*28*). Sequences detected as contained in bacterial chromosomes were removed from further analysis. Three sequences originally identified as plasmids because they cover 80% or the entirety of plasmid CP055640.1 (unnamed plasmid from the *E. coli* strain RHB36-C13) were also removed because they are contained in *E. coli* chromosomes (fig. S14).

#### Mob_recon

The sequences were analysed using mob_recon from the mob_suite v3.0.1 package (*25*) for plasmid reconstruction. We used the options “-u” to “check for circularity flag generated by unicycler”, “-c” to “detect circular contigs with assembly overhangs”, and otherwise default settings.

The initial set of plasmids identified by the *mapping* method (n=408) was combined with those identified by *mob_recon* (n=710), resulting in 765 plasmid sequences (353 identified by both approaches, 357 by *mob_recon* only, and 55 detected exclusively by *mapping*).

### Integrated plasmid dataset

To identify relatives of Murray plasmids, we first generated a comprehensive dataset integrating plasmid sequences from multiple publicly available sources. We started by retrieving sequences from five different plasmid databases commonly cited in plasmid studies: PLSDB v2020_11_19 (*29*), Orlek (*30*), COMPASS (*16*), pATLAS (*31*) and MOB-SUITE v3.0.1 (*25*). To remove redundancy, both the accession numbers and sequences were deduplicated. Sequence duplicates were identified and removed with Dedupe from the BBTools package v38.90 (*32*). The script was run with the flag “ac=f” to remove exact duplicates only and otherwise default parameters. The deduplication step resulted in a core subset of 30,463 sequences (61.05% redundancy) that were subsequently supplemented with sequences from 4 selected plasmid studies (Shaw et al. (*33*), Abe et al. (*34*), Li et al. (*35*) and Leon-Sampedro et al. (*36*)) and new sequences from the latest version of PLSDB (v2021_06_23). The addition of these entries generated a dataset comprising 40,757 plasmid sequences. The total number of plasmid sequences retrieved from each database/dataset is indicated in Fig. 1. The accession numbers of all plasmids in our integrated dataset are provided in table S1.

When available, metadata including host BioSample accession, taxonomy, location, collection date and source of isolation, were retrieved from NCBI for all plasmid sequences available in GenBank/RefSeq using the PLSDB pipeline to process plasmids from NCBI (*29*) (https://github.com/CCB-SB/plsdb/tree/20f77c9bfdfdb82bbacd0085f53b3af98074188a). The metadata linked to the 40,757 plasmid sequences are indicated in table S1.

### Plasmids sequence characterisation

The plasmid sequences in our integrated dataset and those identified in the Murray collection, totalling 41,522, were typed and annotated using a variety of tools and databases (see Fig. 1C). Plasmid mobility classification, relaxase, oriT and MPF typing were performed using mob_typer v3.0.1 (*25*) with default options. Replicon typing was carried out with mob_typer and abricate v1.0.1 (*24*) using the plasmidfinder database (*10*) and a minimum sequence identity (--minid) and coverage (--mincov) of 80 and 60%, respectively.

To comprehensively identify AMR genes, we searched for homologs against multiple databases: CARD (*37*), NCBI (Bioproject PRJNA313047), ARG-ANNOT (*38*) and Resfinder (*39*). To search the different databases in a single step, we integrated them into a single non-redundant database with CD-HIT v4.8.1 (*40*). We used the cd-hit-est algorithm to cluster nucleotide datasets with the following settings: sequence identity threshold of 100% (-c 1.00), word size 10 (-n 10; suggested for thresholds 0.90 ∼ 1.0), and length difference cutoff of 90% (-s 0.9). The databases VFDB (*41*) and Ecoli_VF (*42*) were also combined with CD-HIT (same settings described above) and then used to identify virulence genes. AMR and virulence genes were detected with abricate v1.0.1 (*24*), restricting matches to a minimum coverage and sequence identity of 80% (--mincov, --minid).

Genes encoding transposases were identified via sequence annotation with prokka v1.14.6 (*43*) using the “Bacterial, Archaeal and Plant Plastid” genetic code (--gcode 11) and otherwise default options. Prokka identifies various genomic features, including protein-coding regions, and assigns a function to the gene product based on similarity to known proteins from different databases in a hierarchical order. The first database is a collection of transposase protein sequences from ISfinder, a dedicated database for bacterial insertion sequences (*44*). The number of transposase genes in each plasmid was determined by recording the number of gene products containing the term “transposase” from the prokka output annotation table.

Integrons were detected using Integron finder v2.0.1 (*45*). The program was run with the local detection option (--local-max) to improve sensitivity, the functional annotation option (--func-annot) to annotate gene cassettes with HMM profiles from the AMRFinderPlus database, and the promoter and attI sites search option (--promoter-attI). Additionally, the topology of the input sequence was adjusted to account for circularised plasmids (--circ option). The three types of elements identified by the program and the topology of the input sequence are indicated in table S1.

### Identification of Murray plasmid relatives

To identify relatives of Murray plasmids in current public databases, we compared the 765 plasmids assembled from Murray genomes against the 40,757 sequences comprising our integrated plasmid dataset using two methods:

#### Mash

K-mer-based sketches of all plasmid sequences were generated with the mash v2.3 (*9*) sketch algorithm. A k-mer size of 14 (*k* = 14) was chosen with otherwise default options. The pairwise mutation distances between Murray plasmid sketches and those from the integrated plasmid dataset were estimated using the mash dist algorithm with a distance threshold of 0.05, as previously reported (*9*), and otherwise default settings.

#### Blast

A nucleotide BLAST database of the sequences in the integrated plasmid dataset was created with makeblastdb (from BLAST v2.10.1+ (*46*)). Murray plasmids were compared against the database using blastn with an E-value cutoff of 1e-05, restricting the output to the best non-redundant High-scoring Segment Pairs per subject sequence (option -subject_besthit). Only matches covering at least 70% of the query Murray plasmid were retained as homologs.

Matches detected with *mash* (n=4,923) and *BLAST* (n=9,099) were combined, resulting in the identification of 9,186 plasmids from databases sharing sequence similarity with Murray plasmids (4,836 detected by both mash and blastn, 87 exclusively detected by mash and 4,263 by blastn only). We considered these plasmids as Murray close relatives.

To determine the level of similarity between all (40,757) plasmids comprising our integrated database and Murray plasmids, we repeated the BLAST comparison at the nucleotide level but without filtering matches below 70% coverage of the Murray plasmid. We then extracted the similarity records (e.g. alignment length, query coverage, evalue, bitscore) of the best match for each plasmid from the integrated database featuring significant (E-value cutoff of 1e-05) hits against Murray plasmids. For this analysis, we use the coverage percentage of a Murray plasmid sequence as the criterion to select the best match and the metric to describe similarity (see Fig. 5A, fig. S11 and table s7). Plasmids covering from 1 to 69% of a Murray plasmid sequence were considered Murray extended relatives, and those with no matches (i.e. 0% coverage) Murray unrelated.

### Network clustering and comparative analysis

To determine the diversity of Murray plasmids and their close relatives, we implemented a two-step approach relying on alignment-free sequence comparison and network clustering. First, we estimated sequence similarity based on shared k-mer content between all plasmids using mash with the parameters described in the section above. Second, we used the mash dist output to perform network analyses in Cytoscape v3.8.1 (*47*). The resulting network, representing the patterns of similarity between the 9,951 plasmid sequences, was clustered at two levels with clusterMaker v2.0 (*48*). At the first level, we used the “Connected components cluster (CC)” algorithm with default values to detect all the disconnected components of the network, i.e. clusters of sequences sharing similarity at the minimum mash distance of 0.05. Next, we applied the “Markov Clustering Algorithm (MCL)” to partition the network into groups of strongly connected nodes. MCL is an unsupervised community detection method that simulates flow throughout the network via random walks between nodes. The concept behind this clustering method is that a random walk will more likely remain visiting nodes of a given cluster rather than crossing clusters due to the greater number of links existing within the cluster compared to the fewer occurring between clusters. Hence, the algorithm is able to detect groups of densely connected nodes where the flow gathers (*49*). The dense communities detected in our network correspond to plasmids sharing high genetic similarity. We chose the MCL algorithm because it has been previously applied to the analysis of complex biological similarity networks (*50*). We used the number of shared hashes between plasmids as the array source for MCL, set the edge weight cut-off with the option “Select Cutoff Heuristically” (it calculates a candidate cutoff using the heuristic proposed by Apeltsin et al. (*51*); 400 in our network), and otherwise applied the default parameters. Finally, we combined the results of the two clustering algorithms to assign every plasmid in the network to a group of closely related sequences, hereafter referred to as close families (CFs). The 336 network communities detected by MCL, 264 singletons (i.e. plasmids that did not connect with any other plasmid in the network) detected with the CC algorithm, and 340 plasmids part of connected component clusters that did not cluster with any MCL community (i.e. outliers) were all identified as distinct CFs (See Fig. 1D for a schematic representation), totalling 940. The CF classification of the Murray and related plasmids is indicated in table S2.

The Murray plasmids and close relatives from the integrated dataset isolated during the same period (i.e. up until 1954; n=22) were categorised as pre-antibiotic era (PAE) plasmids; the rest were classified as modern (the isolation date was available for 5,790 out of 9,142 from the corresponding metadata; table S1).

Containment of Murray plasmids within Modern-CF ones was identified from the output of our initial BLAST comparison (i.e. restricted to >=70% Murray plasmid coverage). For every Modern-CF plasmid (n=3,713), we retrieved all matches (n=21,143) against a Murray plasmid and calculated the size ratio of the modern plasmid versus the Murray relative. We retrieved at least one match for 98% (n=3,646) of the Modern-CF plasmids. Next, we parsed the output data to generate a table summarising all the matches between Modern-CF and their contained Murray plasmids along with the corresponding size ratio. Given that a Modern-CF plasmid can match multiple Murray plasmids, we calculated the size ratio using the Modern-CF plasmid size and the average size of the contained Murray plasmids. The resulting table (table S4), which also includes the CF and CF type of the plasmids, was used to generate a network in Cytoscape in which the nodes represent CFs and the connecting edges indicate containment of Murray plasmids and the average size ratio of the containment events. The full network describes 1566 containment interactions between 856 CFs. To improve sparseness and facilitate the visualisation of the network, only interactions featuring a minimum average size ratio of 5 (878 interactions between 526 CFs) are shown in Fig. 2B. The pangenome of CFs comprising more than one plasmid (n=335) was determined with Panaroo v1.2.9 (*52*). We used the prokka annotations as input and ran the program with a sequence identity threshold (--threshold) of 0.7, length difference cutoff (--len_dif_percent) of 0.7, in sensitive stringency mode (--clean-mode), removing invalid annotations (--remove-invalid-genes), and with no edge filtering in the final output graph (--no_clean_edges). The pangenome metrics (table S3) for all CFs were collated from the “summary_statistics.txt” output files. The “core” and “soft_core” components were combined and considered as the core genome (i.e. present in at least 95% of the genomes); “shell” and “cloud” were merged and considered the accessory genome. The average proportion of core genes in a plasmid (see fig. S3) was determined by first calculating the average number of CDS per plasmid in a given CF and then the fraction of them being core according to the pangenome metrics for the CF. The pairwise comparisons of randomly chosen genomes within a plasmid CF were carried out with clinker v0.0.28 (*53*) using prokka annotations as input files and with default settings.

To determine the diversity of the plasmids in the whole integrated database, their relationship to Murray plasmids and their contribution to AMR, we first compared the 40,757 plasmids in the database in an all-vs-all manner using mash with the parameters described in the section above. We then visualised the resulting similarity patterns in a network using Cytoscape, and overlaid either the Murray plasmid coverage values determined by our second BLAST comparison (the one unrestricted to a minimum Murray plasmid coverage) or the number of AMR genes identified in the plasmids (Fig. 5 and table S7).

### Statistical analyses and visualisation

To account for differences in plasmid number across CFs, we used mean values per CF when plotting the distribution of features such as plasmid size, number of replicons, core genes, integrons, and genes encoding AMR determinants, transposases or virulence factors. When relevant, the mean values were further split by mobility type, and by PAE and modern plasmids in Mixed-CFs (e.g. Fig. 3B, fig. S2 and fig. S6).

We used mean values to visually explore size differences between PAE and modern plasmids of the same Mixed-CF and represented them as size ratios in Fig. 4B. To assess the statistical significance of PAE vs modern size differences, we first calculated size differences in PAE vs PAE, modern vs modern, and PAE vs modern plasmids within the same CF. We then compared the resulting distributions with the non-parametric Kolmogorov–Smirnov (K-S) test in a pairwise manner to assess the likelihood of them coming from the same underlying distribution. To account for PAE and modern sampling bias, we implemented a random sampling with replacement strategy when calculating size differences between plasmids of the same CF. In summary, for each of the 88 Mixed-CFs we: 1) randomly sampled with replacement two hundred PAE and modern plasmids (one hundred from each category) and stored their sizes in separate vectors, 2) used the vectors to calculate PAE vs PAE, modern vs modern, and PAE vs modern absolute differences, and 3) performed a K-S test for each comparison (see table S8). A maximum p-value of 0.05 was used to determine if the compared distributions were statistically different. We then plotted the resulting size difference distributions for all CFs (three representative plots are shown in fig. S10 and the whole set is available in the GitHub repository). The same strategy was implemented to assess differences in the number of integrons and genes encoding transposases or virulence factors between PAE and modern plasmids of the same CF (table S8).

Sampling with replacement coupled with a Kolmogorov-Smirnov test was also used to assess the significance of differences between Mixed-CFs with (AMR+; n=39) and without (AMR-; n=49) resistance genes for a number of properties, namely size, core genome, transposases and virulence genes. For these comparisons, mean values per CF were randomly sampled from the AMR+ and AMR-groups (one hundred values each), and the distributions of the sampled values were then compared with the K-S test.

All the plots presented in this study were generated in RStudio v2023.06.0 using R v4.3.1. Networks were generated with Cytoscape v3.8.1. Fig. 1, fig. S4 and fig. S14 were created with BioRender.com. When required, manual figure editing was carried out using Inkscape v1.2.2 (732a01da63, 2022-12-09, custom).

## Acknowledgements

We thank Mathew A. Beale for the helpful discussions during this study.

## Author contributions

A.C., Z.I., and N.R.T. conceived and designed the study. A.C. performed the computational analyses, with contributions from W.F.. W.F. and D.C. contributed to data visualisation. D.C., L.L., J.K., H.M., J.D., and S.A. contributed to data collection. Z.I., and N.R.T supervised the study. A.C., Z.I., W.F., and N.R.T. interpreted the results. A.C., Z.I., and N.R.T. wrote the manuscript, with input from the other authors.

## Data and materials availability

All data are available in the main text or the supplementary materials. The Murray genome sequences are publicly accessible from the BioProject PRJEB3255. Table S1 lists the SRA accessions of the isolates in which plasmids were identified in this study. Our integrated plasmid dataset is comprised of publicly available sequences. Table S1 lists the database from which the plasmids were retrieved, their GenBank accessions, and BioSample accessions when available. Analysis and plotting code is deposited in the GitHub repository https://github.com/biophage/Murray_plasmids.

## Supplementary Materials

***Figs. S1 to S14***

Fig. S1. Plasmids identified in genomes of the Murray collection.

Fig. S2. Characteristics of PAE plasmids and their modern relatives.

Fig. S3. CFs’ pangenomes.

Fig. S4. Distribution of PAE plasmids and their modern close relatives across CF types.

Fig. S5. Murray plasmids containment within Modern-CF plasmids.

Fig. S6. Replicons identified in PAE plasmids and their modern close relatives.

Figure S7. Pairwise comparisons of PAE plasmids and their modern relatives.

Figure S8. Contribution of PAE-related plasmids to AMR.

Figure S9. Distribution of plasmid mobility types across CFs.

Figure S10. Differences between PAE and Modern plasmids’ average size within mixed CFs.

Figure S11. Relationship of Murray plasmids to all sequences from the integrated database.

Figure S12. Comparison of characteristics between plasmids related and unrelated to Murray plasmids.

Figure S13. Characterisation and classification of plasmid sequences from Murray genomes.

Figure S14. Examples of putative Murray plasmid sequences identified as contained in bacterial chromosomes.

***Tables S1 to S8***

Table S1. Characteristics of Murray plasmids and their modern close relatives.

Table S2. Characteristics of plasmid Close Families.

Table S3. Pangenomes of plasmid Close Families.

Table S4. Modern-CF plasmids containing Murray plasmid sequences.

Table S5. Antimicrobial resistance genes identified in Murray plasmid close relatives.

Table S6. Virulence genes identified in Murray plasmid close relatives.

Table S7. Plasmids comprising our integrated dataset and their relationship to Murray plasmids.

Table S8. *P* values of PAE vs Modern and AMR+ vs AMR-comparisons.

## References and Notes

1. C. J. L. Murray, K. S. Ikuta, F. Sharara, L. Swetschinski, G. R. Aguilar, A. Gray, C. Han, C. Bisignano, P. Rao, E. Wool, S. C. Johnson, A. J. Browne, M. G. Chipeta, F. Fell, S. Hackett, G. Haines-Woodhouse, B. H. Kashef Hamadani, E. A. P. Kumaran, B. McManigal, S. Achalapong, R. Agarwal, S. Akech, S. Albertson, J. Amuasi, J. Andrews, A. Aravkin, E. Ashley, F.-X. Babin, F. Bailey, S. Baker, B. Basnyat, A. Bekker, R. Bender, J. A. Berkley, A. Bethou, J. Bielicki, S. Boonkasidecha, J. Bukosia, C. Carvalheiro, C. Castañeda-Orjuela, V. Chansamouth, S. Chaurasia, S. Chiurchiù, F. Chowdhury, R. C. Donatien, A. J. Cook, B. Cooper, T. R. Cressey, E. Criollo-Mora, M. Cunningham, S. Darboe, N. P. J. Day, M. De Luca, K. Dokova, A. Dramowski, S. J. Dunachie, T. D. Bich, T. Eckmanns, D. Eibach, A. Emami, N. Feasey, N. Fisher-Pearson, K. Forrest, C. Garcia, D. Garrett, P. Gastmeier, A. Z. Giref, R. C. Greer, V. Gupta, S. Haller, A. Haselbeck, S. I. Hay, M. Holm, S. Hopkins, Y. Hsia, K. C. Iregbu, J. Jacobs, D. Jarovsky, F. Javanmardi, A. W. J. Jenney, M. Khorana, S. Khusuwan, N. Kissoon, E. Kobeissi, T. Kostyanev, F. Krapp, R. Krumkamp, A. Kumar, H. H. Kyu, C. Lim, K. Lim, D. Limmathurotsakul, M. J. Loftus, M. Lunn, J. Ma, A. Manoharan, F. Marks, J. May, M. Mayxay, N. Mturi, T. Munera-Huertas, P. Musicha, L. A. Musila, M. M. Mussi-Pinhata, R. N. Naidu, T. Nakamura, R. Nanavati, S. Nangia, P. Newton, C. Ngoun, A. Novotney, D. Nwakanma, C. W. Obiero, T. J. Ochoa, A. Olivas-Martinez, P. Olliaro, E. Ooko, E. Ortiz-Brizuela, P. Ounchanum, G. D. Pak, J. L. Paredes, A. Y. Peleg, C. Perrone, T. Phe, K. Phommasone, N. Plakkal, A. Ponce-de-Leon, M. Raad, T. Ramdin, S. Rattanavong, A. Riddell, T. Roberts, J. V. Robotham, A. Roca, V. D. Rosenthal, K. E. Rudd, N. Russell, H. S. Sader, W. Saengchan, J. Schnall, J. A. G. Scott, S. Seekaew, M. Sharland, M. Shivamallappa, J. Sifuentes-Osornio, A. J. Simpson, N. Steenkeste, A. J. Stewardson, T. Stoeva, N. Tasak, A. Thaiprakong, G. Thwaites, C. Tigoi, C. Turner, P. Turner, H. R. van Doorn, S. Velaphi, A. Vongpradith, M. Vongsouvath, H. Vu, T. Walsh, J. L. Walson, S. Waner, T. Wangrangsimakul, P. Wannapinij, T. Wozniak, T. E. M. Young Sharma, K. C. Yu, P. Zheng, B. Sartorius, A. D. Lopez, A. Stergachis, C. Moore, C. Dolecek, M. Naghavi, Global burden of bacterial antimicrobial resistance in 2019: a systematic analysis. Lancet 399, 629–655 (2022).

2. K. C. Nicolaou, S. Rigol, A brief history of antibiotics and select advances in their synthesis. J. Antibiot. 71, 153–184 (2017).

3. S. R. Partridge, S. M. Kwong, N. Firth, S. O. Jensen, Mobile Genetic Elements Associated with Antimicrobial Resistance. Clin. Microbiol. Rev., doi: 10.1128/cmr.00088-17 (2018).

4. S. Castañeda-Barba, E. M. Top, T. Stalder, Plasmids, a molecular cornerstone of antimicrobial resistance in the One Health era. Nat. Rev. Microbiol. 22, 18–32 (2023).

5. J. Davies, D. Davies, Origins and Evolution of Antibiotic Resistance. Microbiol. Mol. Biol. Rev., doi: 10.1128/mmbr.00016-10 (2010).

6. N. Datta, V. M. Hughes, Plasmids of the same Inc groups in Enterobacteria before and after the medical use of antibiotics. Nature 306, 616–617 (1983).

7. K. S. Baker, E. Burnett, H. McGregor, A. Deheer-Graham, C. Boinett, G. C. Langridge, M. Wailan, A. K. Cain, N. R. Thomson, J. E. Russell, J. Parkhill, The Murray collection of pre-antibiotic era Enterobacteriacae: a unique research resource. Genome Med. 7, 1–7 (2015).

8. World Health Organization, WHO Bacterial Priority Pathogens List, 2024:Bacterial Pathogens of Public Health Importance, to Guide Research, Development, and Strategies to Prevent and Control Antimicrobial Resistance (World Health Organization, 2024).

9. B. D. Ondov, T. J. Treangen, P. Melsted, A. B. Mallonee, N. H. Bergman, S. Koren, A. M. Phillippy, Mash: fast genome and metagenome distance estimation using MinHash. Genome Biol. 17, 1–14 (2016).

10. A. Carattoli, E. Zankari, A. García-Fernández, M. V. Larsen, O. Lund, L. Villa, F. M. Aarestrup, H. Hasman, In Silico Detection and Typing of Plasmids using PlasmidFinder and Plasmid Multilocus Sequence Typing. Antimicrob. Agents Chemother., doi: 10.1128/aac.02412-14 (2014).

11. M. Castanheira, P. J. Simner, P. A. Bradford, Extended-spectrum β-lactamases: an update on their characteristics, epidemiology and detection. JAC Antimicrob Resist 3, dlab092 (2021).

12. C. Smillie, M. Pilar Garcillán-Barcia, M. Victoria Francia, E. P. C. Rocha, F. de la Cruz, Mobility of Plasmids. Microbiol. Mol. Biol. Rev., doi: 10.1128/mmbr.00020-10 (2010).

13. K. Hardy, Bacterial Plasmids (Springer Science & Business Media, 1986).

14. V. M. Hughes, N. Datta, Conjugative plasmids in bacteria of the “pre-antibiotic” era. Nature 302, 725–726 (1983).

15. Plasmid cointegrates and their resolution mediated by transposon Tn3 mutants. Gene 15, 103–118 (1981).

16. P.-E. Douarre, L. Mallet, N. Radomski, A. Felten, M.-Y. Mistou, Analysis of COMPASS, a New Comprehensive Plasmid Database Revealed Prevalence of Multireplicon and Extensive Diversity of IncF Plasmids. Front. Microbiol. 11, 503503 (2020).

17. R. R. Wick, L. M. Judd, C. L. Gorrie, K. E. Holt, Unicycler: Resolving bacterial genome assemblies from short and long sequencing reads. PLoS Comput. Biol. 13, e1005595 (2017).

18. A. Gurevich, V. Saveliev, N. Vyahhi, G. Tesler, QUAST: quality assessment tool for genome assemblies. Bioinformatics 29, 1072–1075 (2013).

19. D. E. Wood, J. Lu, B. Langmead, Improved metagenomic analysis with Kraken 2. Genome Biol. 20, 1–13 (2019).

20. J. Lu, F. P. Breitwieser, P. Thielen, S. L. Salzberg, Bracken: estimating species abundance in metagenomics data. PeerJ Comput. Sci. 3, e104 (2017).

21. D. Antipov, M. Raiko, A. Lapidus, P. A. Pevzner, Plasmid detection and assembly in genomic and metagenomic data sets. Genome Res. 29, 961–968 (2019).

22. S. R. Eddy, Accelerated Profile HMM Searches. PLoS Comput. Biol. 7, e1002195 (2011).

23. M. Hunt, N. D. Silva, T. D. Otto, J. Parkhill, J. A. Keane, S. R. Harris, Circlator: automated circularization of genome assemblies using long sequencing reads. Genome Biol. 16, 294 (2015).

24. GitHub - tseemann/abricate: :mag_right: Mass screening of contigs for antimicrobial and virulence genes, GitHub. https://github.com/tseemann/abricate.

25. J. Robertson, J. H. E. Nash, MOB-suite: software tools for clustering, reconstruction and typing of plasmids from draft assemblies. Microb Genom 4 (2018).

26. H. Li, Minimap2: pairwise alignment for nucleotide sequences. Bioinformatics 34, 3094–3100 (2018).

27. GitHub - mbhall88/pafpy: A lightweight library for working with PAF (Pairwise mApping Format) files, GitHub. https://github.com/mbhall88/pafpy.

28. M. Johnson, I. Zaretskaya, Y. Raytselis, Y. Merezhuk, S. McGinnis, T. L. Madden, NCBI BLAST: a better web interface. Nucleic Acids Res. 36, W5–W9 (2008).

29. V. Galata, T. Fehlmann, C. Backes, A. Keller, PLSDB: a resource of complete bacterial plasmids. Nucleic Acids Res. 47, D195–D202 (2019).

30. A curated dataset of complete Enterobacteriaceae plasmids compiled from the NCBI nucleotide database. Data in Brief 12, 423–426 (2017).

31. T. F. Jesus, B. Ribeiro-Gonçalves, D. N. Silva, V. Bortolaia, M. Ramirez, J. A. Carriço, Plasmid ATLAS: plasmid visual analytics and identification in high-throughput sequencing data. Nucleic Acids Res. 47, D188–D194 (2018).

32. BBMap, SourceForge (2022). https://sourceforge.net/projects/bbmap/.

33. L. P. Shaw, K. K. Chau, J. Kavanagh, M. AbuOun, E. Stubberfield, H. Soon Gweon, L. Barker, G. Rodger, M. J. Bowes, A. T. M. Hubbard, H. Pickford, J. Swann, D. Gilson, R.P. Smith, S. J. Hoosdally, R. Sebra, H. Brett, T. E. A. Peto, M. J. Bailey, D. W. Crook, D. S. Read, M. F. Anjum, A. Sarah Walker, N. Stoesser, REHAB consortium, Niche and local geography shape the pangenome of wastewater- and livestock-associated Enterobacteriaceae. Science Advances, doi: 10.1126/sciadv.abe3868 (2021).

34. R. Abe, Y. Akeda, Y. Sugawara, D. Takeuchi, Y. Matsumoto, D. Motooka, N. Yamamoto, R. Kawahara, K. Tomono, Y. Fujino, S. Hamada, Characterization of the Plasmidome Encoding Carbapenemase and Mechanisms for Dissemination of Carbapenem-Resistant Enterobacteriaceae. mSystems, doi: 10.1128/msystems.00759-20 (2020).

35. R. Li, X. Lu, K. Peng, Z. Liu, Y. Li, Y. Liu, X. Xiao, Z. Wang, Deciphering the Structural Diversity and Classification of the Mobile Tigecycline Resistance Gene tet(X)-Bearing Plasmidome among Bacteria. mSystems, doi: 10.1128/msystems.00134-20 (2020).

36. R. León-Sampedro, J. DelaFuente, C. Díaz-Agero, T. Crellen, P. Musicha, J. Rodríguez-Beltrán, C. de la Vega, M. Hernández-García, N. López-Fresneña, P. Ruiz-Garbajosa, R. Cantón, B. S. Cooper, Á. San Millán, Pervasive transmission of a carbapenem resistance plasmid in the gut microbiota of hospitalized patients. Nature Microbiology 6, 606–616 (2021).

37. B. P. Alcock, A. R. Raphenya, T. T. Y. Lau, K. K. Tsang, M. Bouchard, A. Edalatmand, W. Huynh, A.-L. V. Nguyen, A. A. Cheng, S. Liu, S. Y. Min, A. Miroshnichenko, H.-K. Tran, R. E. Werfalli, J. A. Nasir, M. Oloni, D. J. Speicher, A. Florescu, B. Singh, M. Faltyn, A. Hernandez-Koutoucheva, A. N. Sharma, E. Bordeleau, A. C. Pawlowski, H. L. Zubyk, D. Dooley, E. Griffiths, F. Maguire, G. L. Winsor, R. G. Beiko, F. S. L. Brinkman, W. W. L. Hsiao, G. V. Domselaar, A. G. McArthur, CARD 2020: antibiotic resistome surveillance with the comprehensive antibiotic resistance database. Nucleic Acids Res. 48, D517–D525 (2020).

38. S. K. Gupta, B. R. Padmanabhan, S. M. Diene, R. Lopez-Rojas, M. Kempf, L. Landraud, J.-M. Rolain, ARG-ANNOT, a new bioinformatic tool to discover antibiotic resistance genes in bacterial genomes. Antimicrob. Agents Chemother. 58, 212–220 (2014).

39. E. Zankari, H. Hasman, S. Cosentino, M. Vestergaard, S. Rasmussen, O. Lund, F. M. Aarestrup, M. V. Larsen, Identification of acquired antimicrobial resistance genes. J. Antimicrob. Chemother. 67, 2640–2644 (2012).

40. W. Li, A. Godzik, Cd-hit: a fast program for clustering and comparing large sets of protein or nucleotide sequences. Bioinformatics 22, 1658–1659 (2006).

41. L. Chen, D. Zheng, B. Liu, J. Yang, Q. Jin, VFDB 2016: hierarchical and refined dataset for big data analysis--10 years on. Nucleic Acids Res. 44, D694–7 (2016).

42. GitHub - phac-nml/ecoli_vf: Curated virulence factors for Escherichia coli, GitHub. https://github.com/phac-nml/ecoli_vf.

43. T. Seemann, Prokka: rapid prokaryotic genome annotation. Bioinformatics 30, 2068–2069 (2014).

44. P. Siguier, J. Perochon, L. Lestrade, J. Mahillon, M. Chandler, ISfinder: the reference centre for bacterial insertion sequences. Nucleic Acids Res. 34, D32–D36 (2006).

45. B. Néron, E. Littner, M. Haudiquet, A. Perrin, J. Cury, E. P. C. Rocha, IntegronFinder 2.0: Identification and Analysis of Integrons across Bacteria, with a Focus on Antibiotic Resistance in Klebsiella. Microorganisms 10, 700 (2022).

46. Basic local alignment search tool. J. Mol. Biol. 215, 403–410 (1990).

47. P. Shannon, A. Markiel, O. Ozier, N. S. Baliga, J. T. Wang, D. Ramage, N. Amin, B. Schwikowski, T. Ideker, Cytoscape: A Software Environment for Integrated Models of Biomolecular Interaction Networks. Genome Res. 13, 2498–2504 (2003).

48. J. H. Morris, L. Apeltsin, A. M. Newman, J. Baumbach, T. Wittkop, G. Su, G. D. Bader, T. E. Ferrin, clusterMaker: a multi-algorithm clustering plugin for Cytoscape. BMC Bioinformatics 12, 1–14 (2011).

49. S. Van Dongen, Graph Clustering Via a Discrete Uncoupling Process. SIAM J. Matrix Anal. Appl., doi: 10.1137/040608635 (2008).

50. A. J. Enright, S. Van Dongen, C. A. Ouzounis, An efficient algorithm for large-scale detection of protein families. Nucleic Acids Res. 30, 1575–1584 (2002).

51. L. Apeltsin, J. H. Morris, P. C. Babbitt, T. E. Ferrin, Improving the quality of protein similarity network clustering algorithms using the network edge weight distribution. Bioinformatics 27 (2011).

52. G. Tonkin-Hill, N. MacAlasdair, C. Ruis, A. Weimann, G. Horesh, J. A. Lees, R. A. Gladstone, S. Lo, C. Beaudoin, R. A. Floto, S. D. W. Frost, J. Corander, S. D. Bentley, J. Parkhill, Producing polished prokaryotic pangenomes with the Panaroo pipeline. Genome Biol. 21, 180 (2020).

53. C. L. M. Gilchrist, Y.-H. Chooi, clinker & clustermap.js: automatic generation of gene cluster comparison figures. Bioinformatics 37, 2473–2475 (2021).

