## Supplementary_figures for "Pre and Post antibiotic epoch: insights into the historical spread of antimicrobial resistance"

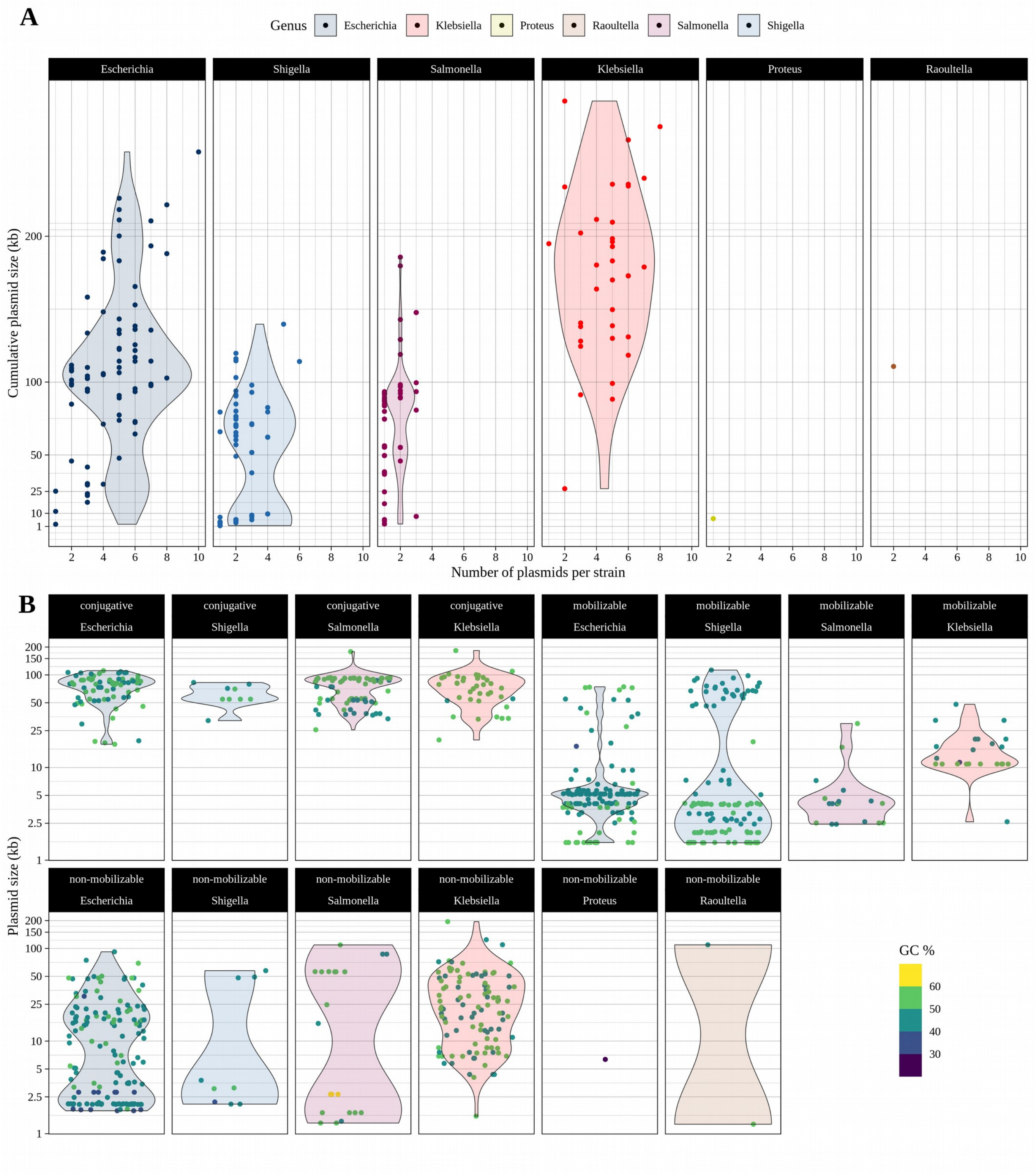


**Fig. S1. Plasmids identified in genomes of the Murray collection.** (**A**) Summary of plasmid content in strains of the Murray collection. Each point in the plots (split by genus) represents a Murray strain and is the function of the number of plasmids identified and the sum of their sizes. (**B**) Distribution of plasmids size. Dots represent Murray plasmids. Plots are split by genus and mobility classification.

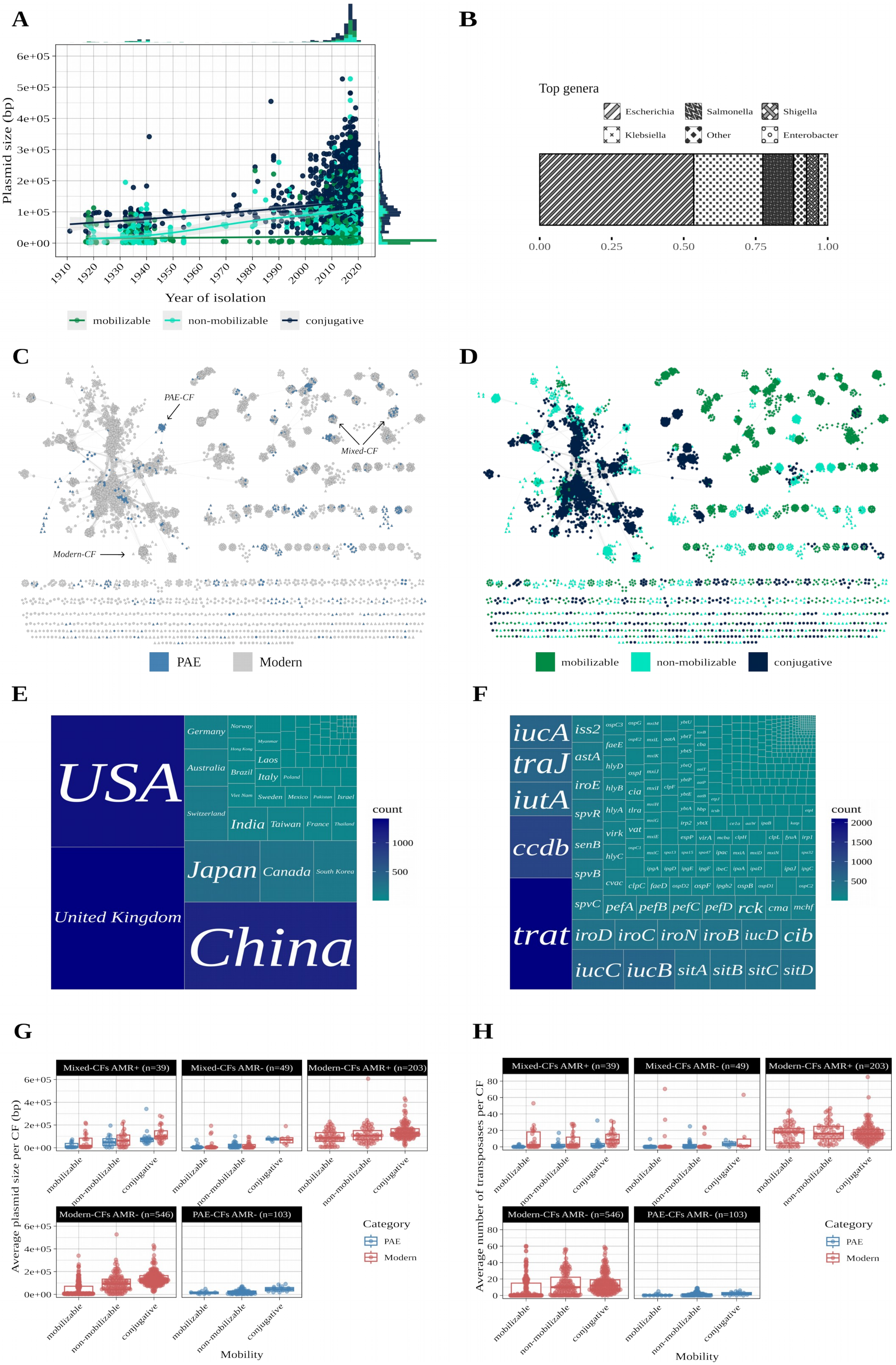

A
Plasmid size (bp)
6e+05
5e+05
4e+05
3e+05
2e+05
1e+05
0e+00
1910
1920
1930
1940
1950
1960
1970
1980
1990
2000
2010
2020
Year of isolation
mobilizable
non-mobilizable
conjugative
B
Top genera
Escherichia
Salmonella
Shigella
Klebsiella
Other
Enterobacter
0.00
0.25
0.50
0.75
1.00
C
PAE-CF
Mixed-CF
Modern-CF
PAE
Modern
D
mobilizable
non-mobilizable
conjugative
E
USA
United Kingdom
China
Japan
Canada
South Korea
India
Taiwan
France
Thailand
Germany
Australia
Switzerland
Brazil
Italy
Laos
Sweden
Mexico
Israel
count
1000
500
F
iucA
traJ
iutA
ccdb
trat
iss2
astA
iroE
spvR
senB
spvB
spvC
hlyD
hlyB
hlyA
virk
hlyC
cvac
pefA
pefB
pefC
pefD
rck
cma
mchf
iroD
iroC
iroN
iroB
iucD
cib
iucC
iucB
sitA
sitB
sitC
sitD
count
2000
1500
1000
500
G
Mixed-CFs AMR+ (n=39)
Mixed-CFs AMR- (n=49)
Modern-CFs AMR+ (n=203)
Modern-CFs AMR- (n=546)
PAE-CFs AMR- (n=103)
Average plasmid size per CF (bp)
6e+05
4e+05
2e+05
0e+00
mobilizable
non-mobilizable
conjugative
Mobility
Category
PAE
Modern
H
Mixed-CFs AMR+ (n=39)
Mixed-CFs AMR- (n=49)
Modern-CFs AMR+ (n=203)
Modern-CFs AMR- (n=546)
PAE-CFs AMR- (n=103)
Average number of transposases per CF
80
60
40
20
0
mobilizable
non-mobilizable
conjugative
Mobility
Category
PAE
Modern

**Fig. S2. Characteristics of PAE plasmids and their modern relatives.** (**A**) Size and year of isolation. Plasmids isolated on or before 1954 were considered PAE in this study. (**B**) Plasmids taxonomic distribution. (**C**) Similarity network illustrating the diversity and distribution of PAE and modern plasmids. Examples of the three types of plasmid CF based on plasmid composition are indicated in the network. (**D**) Similarity network illustrating the distribution of plasmid mobility types. (**E**) Geographical distribution of the plasmids host. (**H**) Diversity and abundance of virulence genes identified in the plasmids. (**G**) Average plasmid size across CFs. (**H**) Average number of transposases across CFs. In panels **G** and **H**, plots are split by CF type. Points represent average values per CF, split by PAE and modern plasmids in Mixed-CFs.

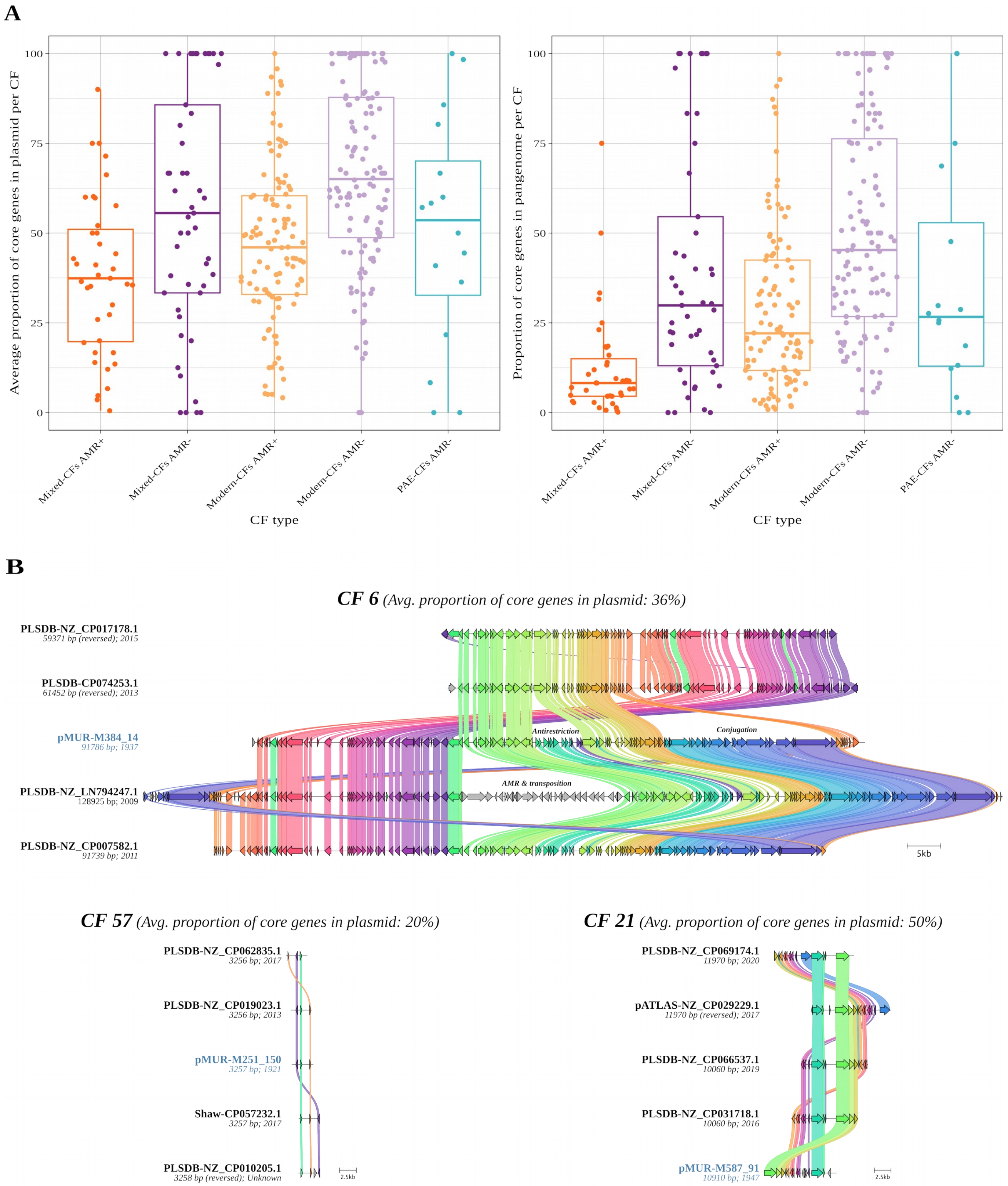


**Fig. S3. CFs' pangenomes.** (**A**) Core component of CF pangenomes. Points in the plots represent the average proportion of genes that are core in a plasmid (left) or the fraction of genes that are core in the pangenome (right) of a given CF. Core genes are those present in at least 95% of the genomes, as determined by Panaroo. Colours denote different CF types, as shown in Fig. 2A. (**B**) Pairwise comparison of plasmids from the same CF. Nucleotide-level comparison of five plasmids of the same CF for three different Mixed-CFs. The selected CFs represent different points in the distribution shown for Mixed-CFs containing resistance plasmids (Mixed-CFs AMR+) in the left plot of panel **A**. The five plasmid sequences were selected randomly. Murray plasmids are highlighted with blue labels. For the comparison of plasmids from CF 6, the functional category of genes located in three indels regarding the Murray plasmid reference is indicated in the figure.

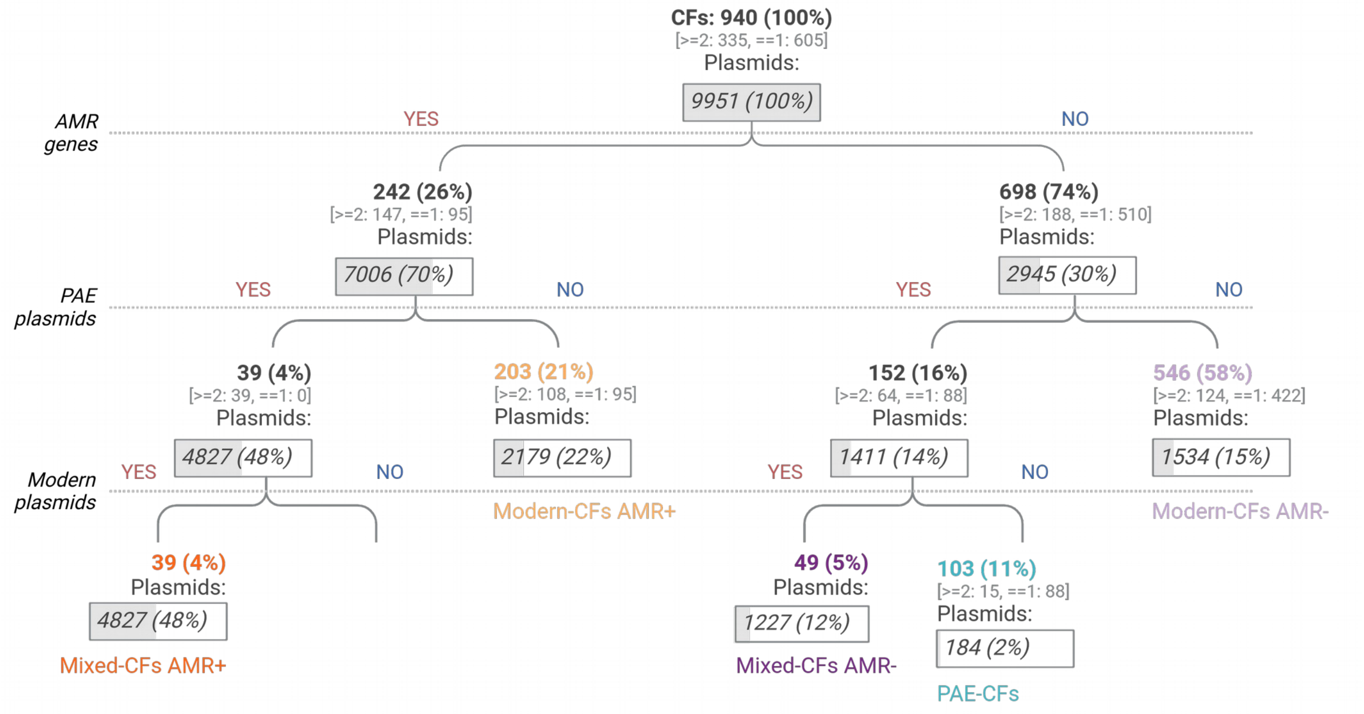


**Fig. S4. Distribution of PAE plasmids and their modern close relatives across CF types.** The hierarchical tree illustrates the classification of plasmid CFs into CF types based on the presence of AMR genes and plasmid composition (presence of PAE and/or modern plasmids). It also shows the breakdown of the number and percentage of CFs and plasmids across the classification steps. The number of CFs containing one (==1) or more than one (>=2) plasmid is indicated in square brackets. The colours of the CF-type labels match those of the network in Fig. 2A.

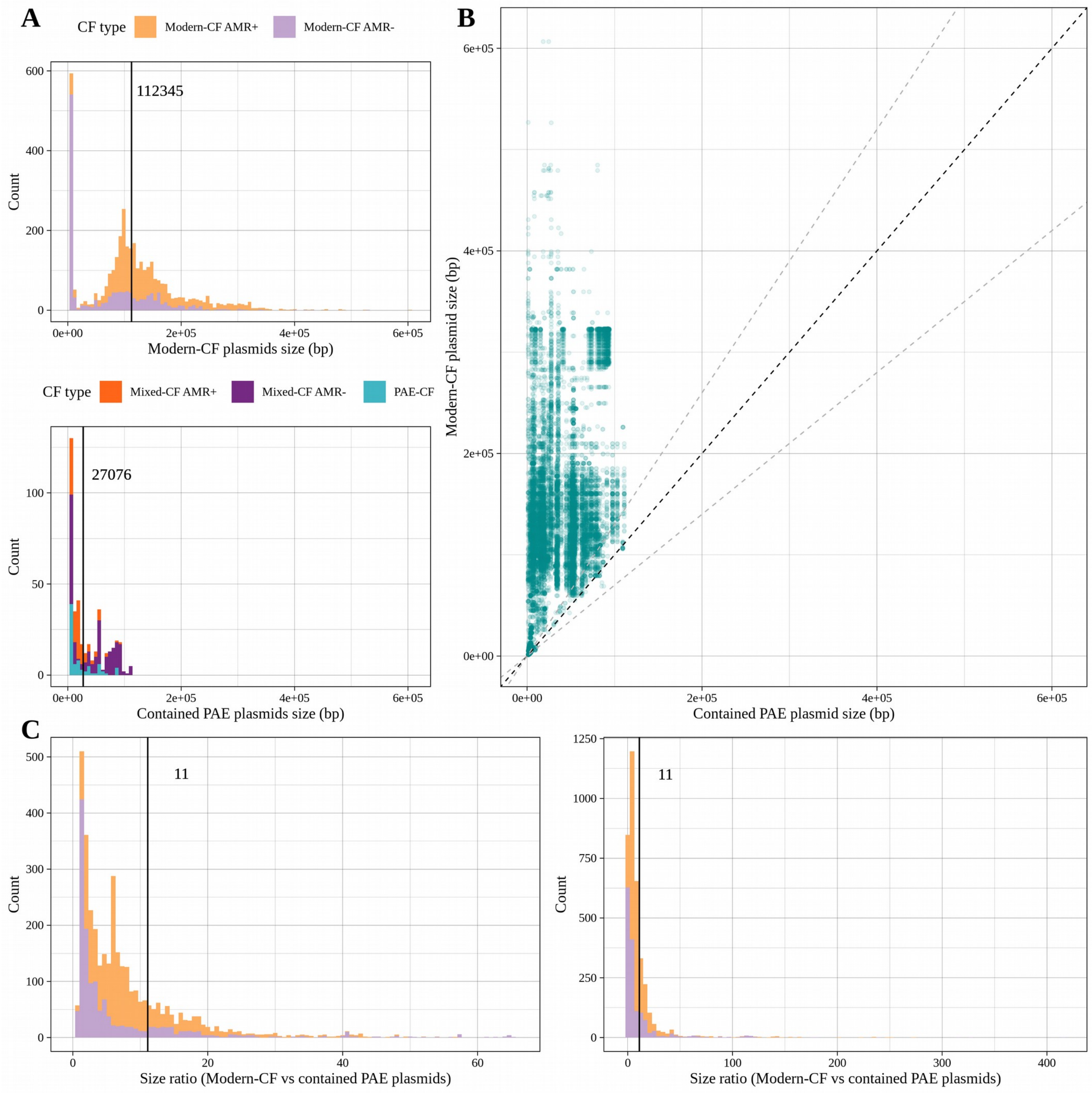


**Fig. S5. Murray plasmids containment within Modern-CF plasmids.** (**A**) Size distribution of Modern-CF plasmids (top) and the Murray plasmids they contain (bottom). The mean size is indicated with a black vertical line in both plots. (**B**) Size relationship between Modern-CF and contained Murray plasmids for all the containment interactions identified. The plot shows the size of Modern-CF and contained Murray plasmids for the 21,143 matches detected via BLAST. Note that a given Modern-CF plasmid can match different Murray plasmids, resulting in multiple points in the plot. The black dotted line indicates a size ratio of 1; the adjacent grey dotted lines correspond to size ratios of 0.7 (below) and 1.3 (above). (**C**) Distribution of Modern-CF vs contained Murray plasmid size ratios. Given that a Modern-CF plasmid can match multiple Murray plasmids, the size ratio shown in the plots represents the difference between the Modern-CF plasmid size and the average size of the contained Murray plasmids. A black vertical line indicates the mean size ratio. The left plot shows the distribution excluding outliers (defined as the values beyond the 0.975 quantile).

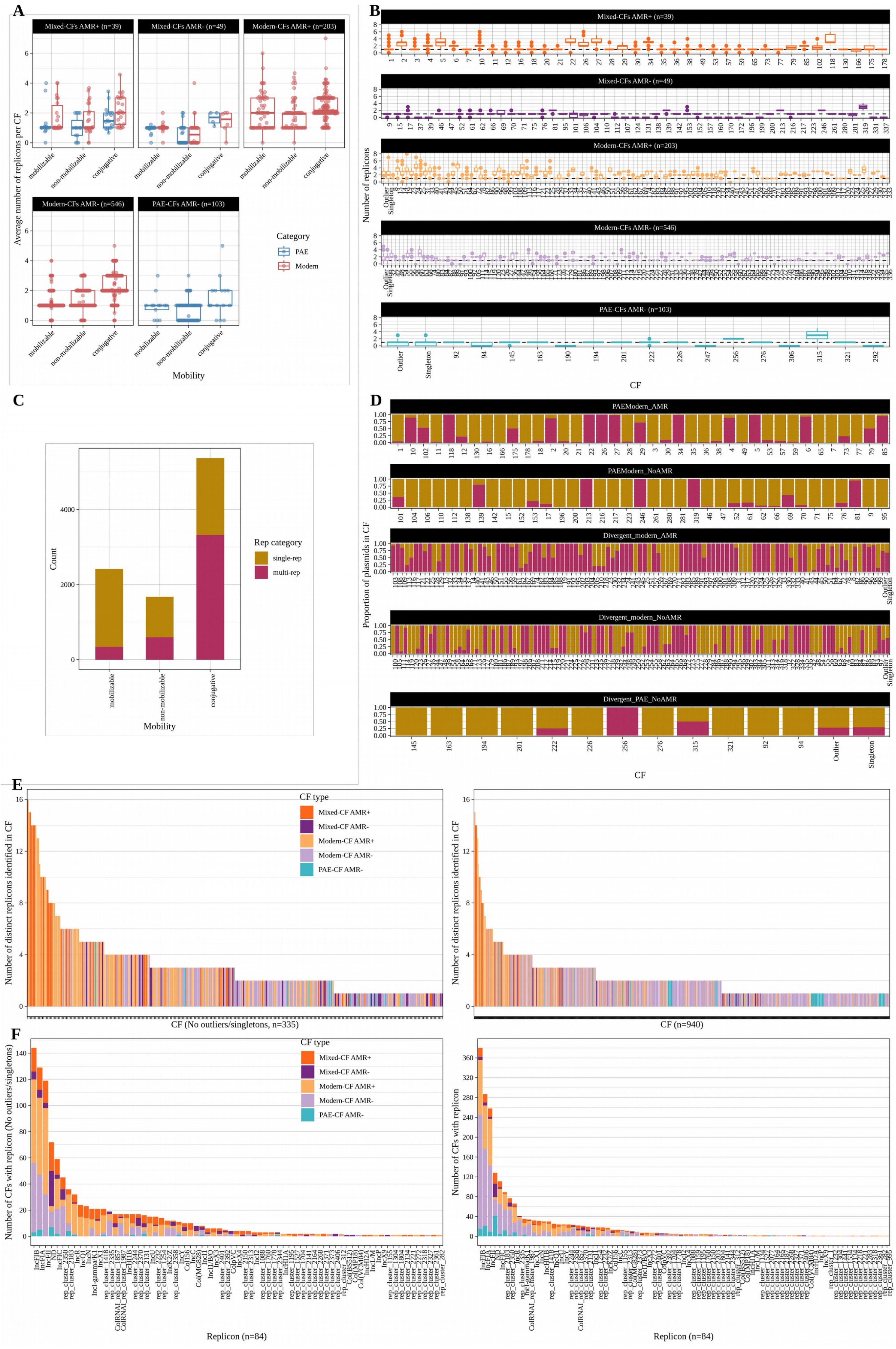

A
Mixed-CFs AMR+ (n=39)
Mixed-CFs AMR- (n=49)
Modern-CFs AMR+ (n=203)
Modern-CFs AMR- (n=546)
PAE-CFs AMR- (n=103)
Average number of replicons per CF
mobilizable
non-mobilizable
conjugative
Mobility
Category
PAE
Modern
B
Mixed-CFs AMR+ (n=39)
Mixed-CFs AMR- (n=49)
Modern-CFs AMR+ (n=203)
Modern-CFs AMR- (n=546)
PAE-CFs AMR- (n=103)
Number of replicons
CF
C
Count
Rep category
single-rep
multi-rep
mobilizable
non-mobilizable
conjugative
Mobility
D
PAEModern_AMR
PAEModern_NoAMR
Divergent_modern_AMR
Divergent_modern_NoAMR
Divergent_PAE_NoAMR
Proportion of plasmids in CF
CF
E
CF type
Mixed-CF AMR+
Mixed-CF AMR-
Modern-CF AMR+
Modern-CF AMR-
PAE-CF AMR-
Number of distinct replicons identified in CF
CF (No outliers/singletons, n=335)
CF (n=940)
F
CF type
Mixed-CF AMR+
Mixed-CF AMR-
Modern-CF AMR+
Modern-CF AMR-
PAE-CF AMR-
Number of CFs with replicon (No outliers/singletons)
Number of CFs with replicon
Replicon (n=84)
Replicon (n=84)

**Fig. S6. Replicons identified in PAE plasmids and their modern close relatives.** (**A**) Average number of replicons identified per plasmid across CFs. The plot is split by CF type. Points represent average values per CF, split by PAE and modern plasmids in Mixed-CFs. (**B**) Distribution of the number of replicons identified in plasmids across CFs. The black horizontal dotted line corresponds to a value of 1. Colours denote CF types as shown in Fig. 2A. (**C**) Proportion of plasmids with single (single-rep) or multiple (multi-rep) replicons across mobility types. (**D**) Proportion of single and muli-replicon plasmids across CFs. (**E**) Number of distinct replicon types identified across CFs. CFs comprising one plasmid are excluded in the left plot. (**F**) Replicons occurrence across CFs. The left plot excludes CFs comprising one plasmid.

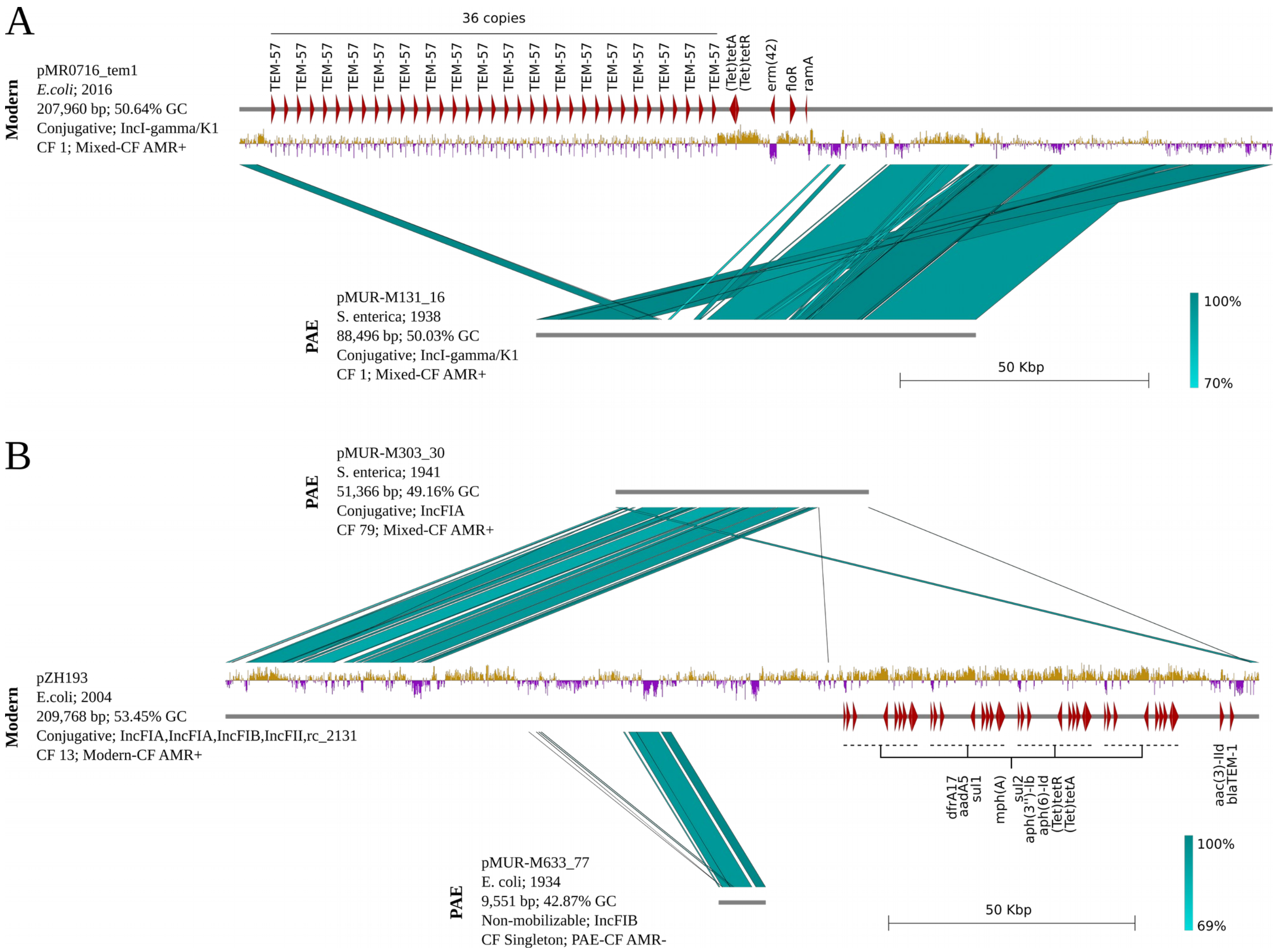


**Figure S7. Pairwise comparisons of PAE plasmids and their modern relatives.** (**A**) An example of a plasmid backbone that maintained a clonal frame during the antibiotic era. (**B**) An example of a modern plasmid (middle) resulting from plasmid fusion, including two PAE plasmid backbones (top and bottom). In both panels, AMR genes encoded in the modern plasmid are depicted as red arrows. GC content distribution is plotted below the grey bar representing the plasmid genome; gold and purple denote above and below the average values, respectively. Tandem duplication of genes (panel **A**) or gene arrays (panel **B**) is indicated above or below the AMR genes.

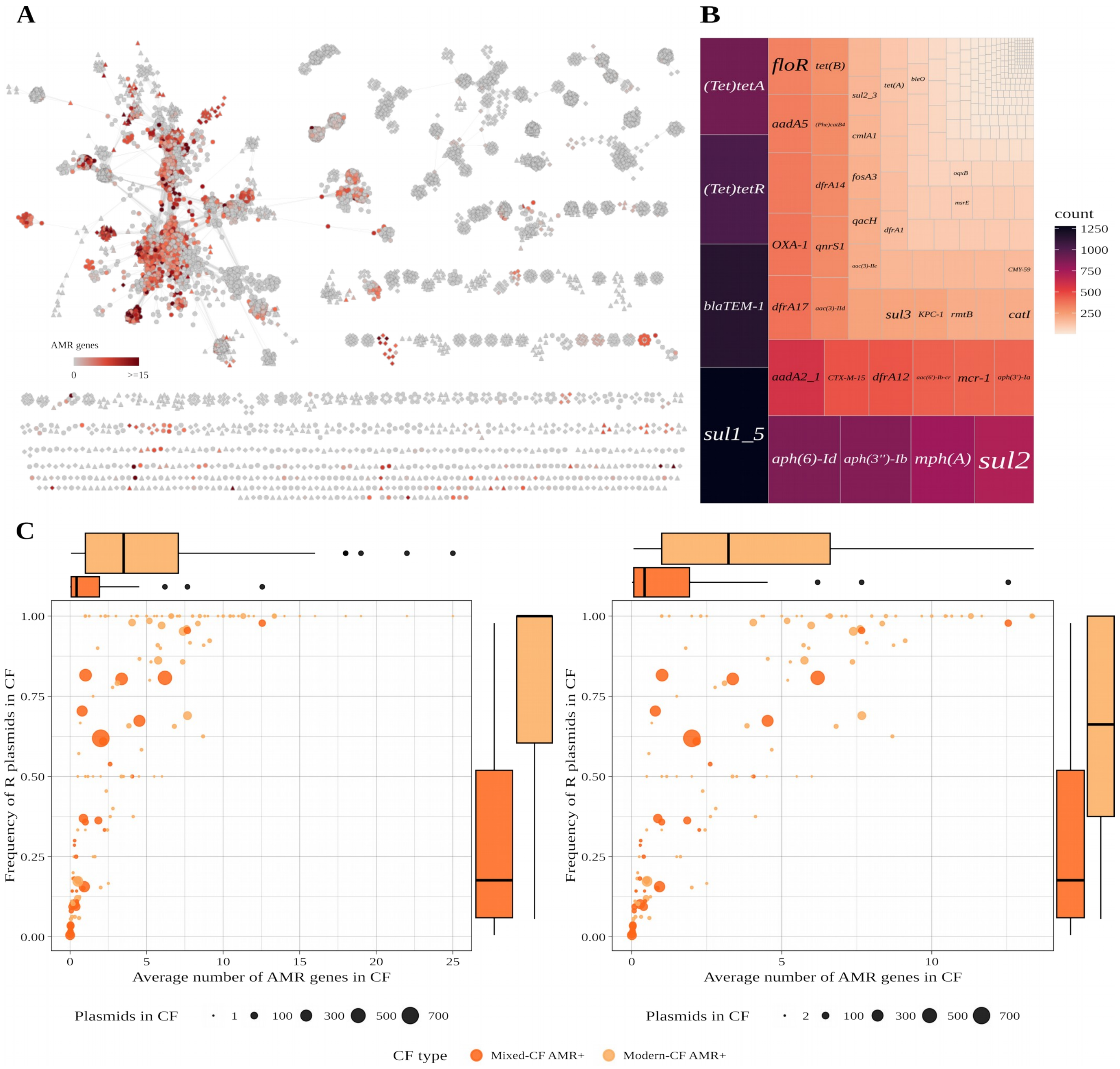


**Figure S8. Contribution of PAE-related plasmids to AMR.** (**A**) Mash similarity network of PAE plasmids and their close relatives illustrating the distribution of resistance plasmids and AMR content. No AMR genes were detected in PAE plasmids. (**B**) Diversity and abundance of AMR genes identified in modern plasmids. (**C**) Contribution of plasmid CFs to AMR. The plots show the average number of AMR genes carried by plasmids (X-axis) and the proportion of resistance plasmids (Y-axis) within a given CF. The right plot excludes CFs comprising one plasmid. The colour of the data points differentiates Mixed- (dark orange) and Modern-CFs (light orange).

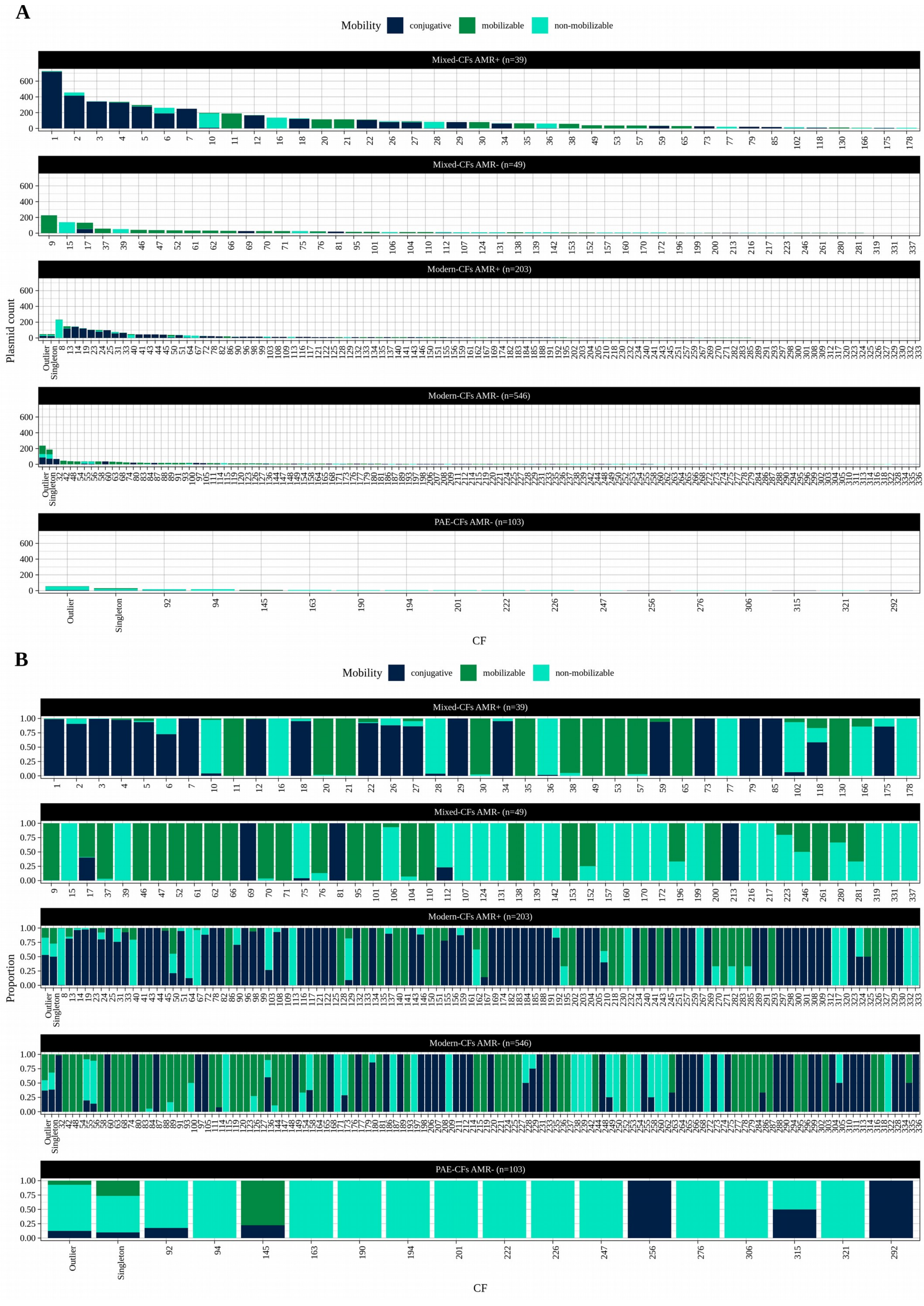


**Figure S9. Distribution of plasmid mobility types across CFs.** (**A**) Number of plasmids within a CF classified by mobility types. (**B**) Proportion of plasmids within a CF classified by mobility types.

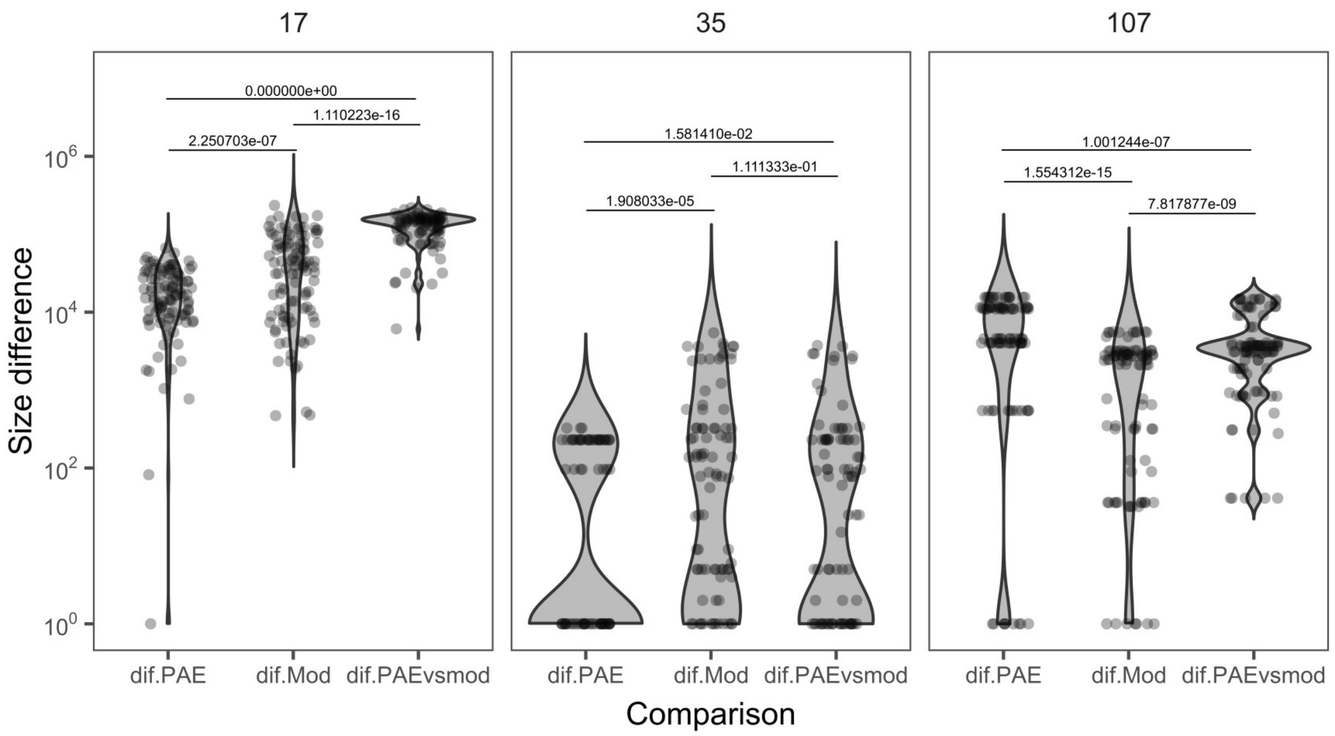


**Figure S10. Differences between PAE and Modern plasmids' average size within mixed CFs.** To assess the significance of PAE vs modern size differences within a Mixed-CF, we used the Kolmogorov–Smirnov (K-S) test to compare the distribution of differences resulting from comparing PAE vs PAE (dif.PAE), modern vs modern (diff.Mod) and PAE vs modern (dif.PAEvsmod) plasmids. Plasmids were selected randomly to account for sampling bias, and the process was repeated 100 times (see Methods). The plots show the distribution of size differences calculated from the randomly picked plasmids. Each data point represents the absolute size difference between a pair of plasmids. The p-value of the K-S test is indicated above the corresponding distributions. The three plots represent exemplars of Mixed-CFs (indicated above the plot) featuring marked (left), moderate (right) or marginal (aka size stasis; middle) size differences between PAE and modern plasmids according to the comparison of their average values (see Fig. 4B). The complete set of plots for the 88 Mixed-CFs is available in the GitHub repository for this study. The same strategy was used to assess the PAE vs Modern differences in cargo genes; the results are provided in table S8.

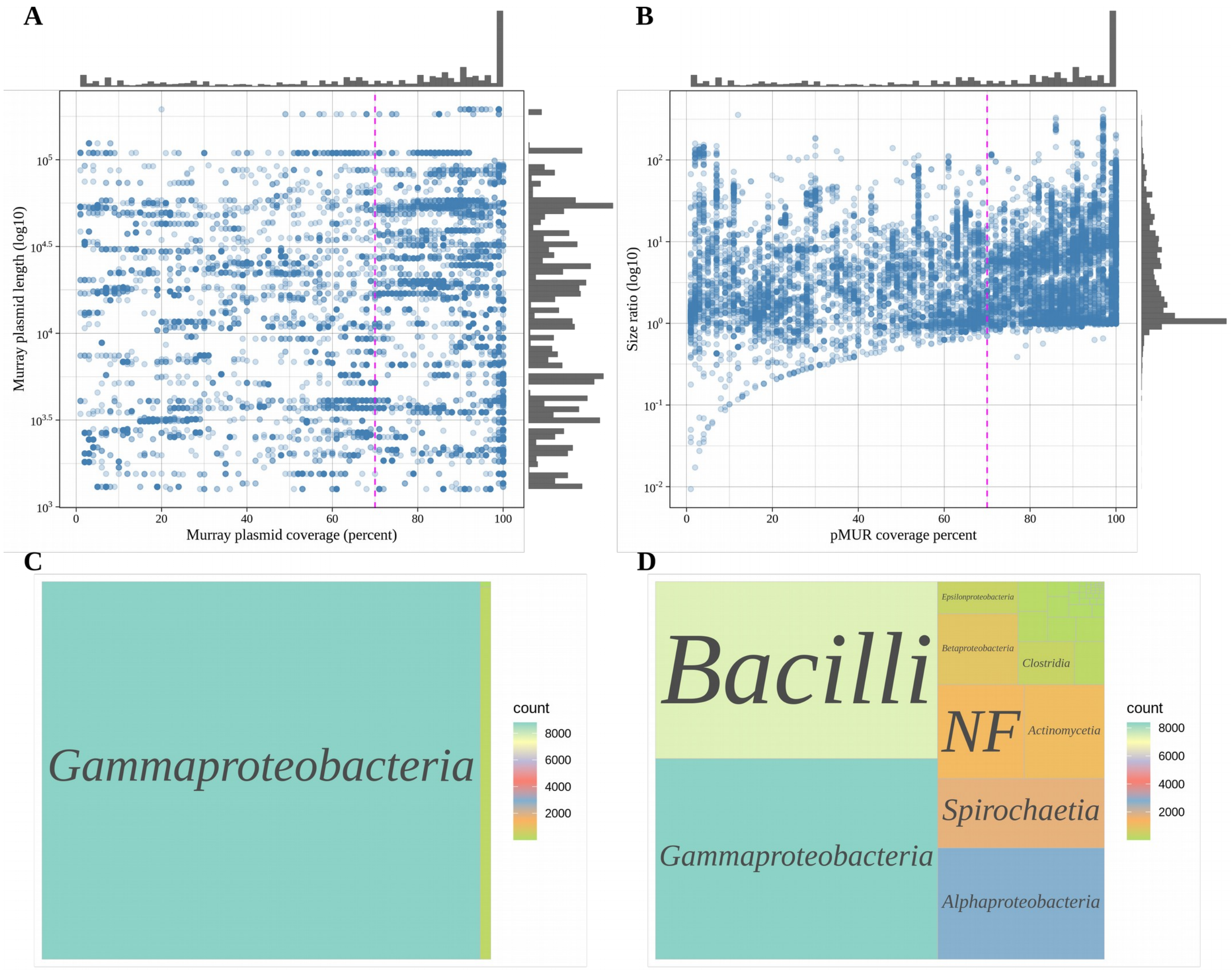


**Figure S11. Relationship of Murray plasmids to all sequences from the integrated database.** (**A**) Distribution of matches between sequences in the integrated plasmid database and Murray plasmids. The plot illustrates the nucleotide-level similarity, expressed as sequence coverage, between 15,644 plasmids from our integrated dataset (see Methods and Fig. 1B) and Murray plasmids. Data points represent the best (highest Murray plasmid coverage) match identified for a given plasmid in the integrated database with a minimum Murray coverage value of 1% detected via BLAST. (**B**) Size ratio between plasmids from the integrated dataset and their closest Murray-plasmid relative for the matches shown in panel A. A vertical pink dotted line in panels A and B marks 70% coverage, one of the criteria used to identify Murray-plasmid close relatives in this study (Fig. 1C). (**C**) Host taxonomic distribution at the level of class for close relatives of Murray plasmids (>= 70% Murray-plasmid coverage). (**D**) Host taxonomic distribution at the level of class for close Murray-unrelated plasmids (0% Murray-plasmid coverage; see Fig. 5A).

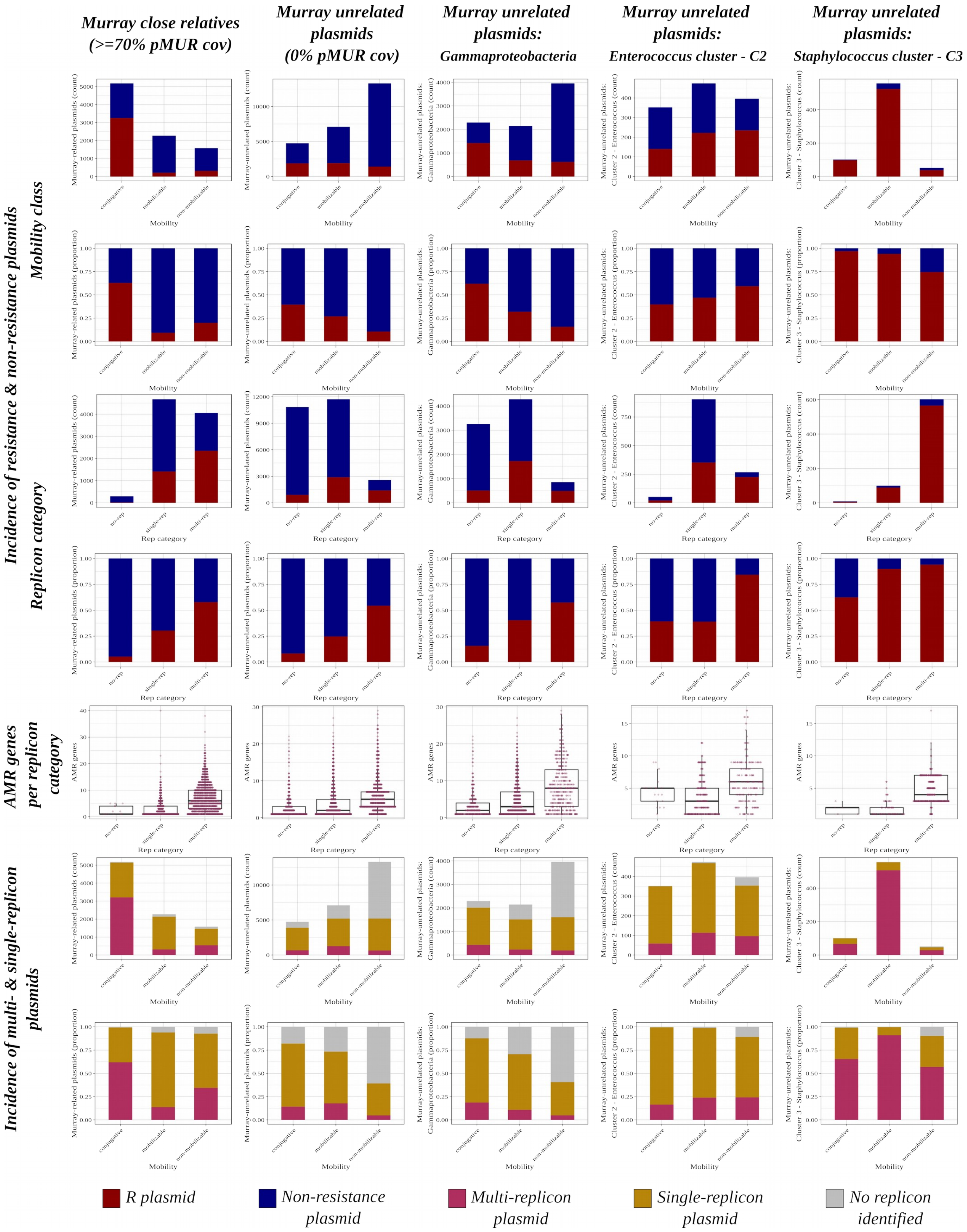


**Figure S12. Comparison of characteristics between plasmids related and unrelated to Murray plasmids.** The figure contrasts the distribution of resistance plasmids (rows 1-4), AMR carriage (row 5), and multi-replicon prevalence (rows 6 and 7) for plasmids from databases closely related to Murray plasmids (column 1) or those unrelated (columns 2-5). Columns 3 to 5 correspond to subsets of Murray-unrelated plasmids as indicated in the column labels.

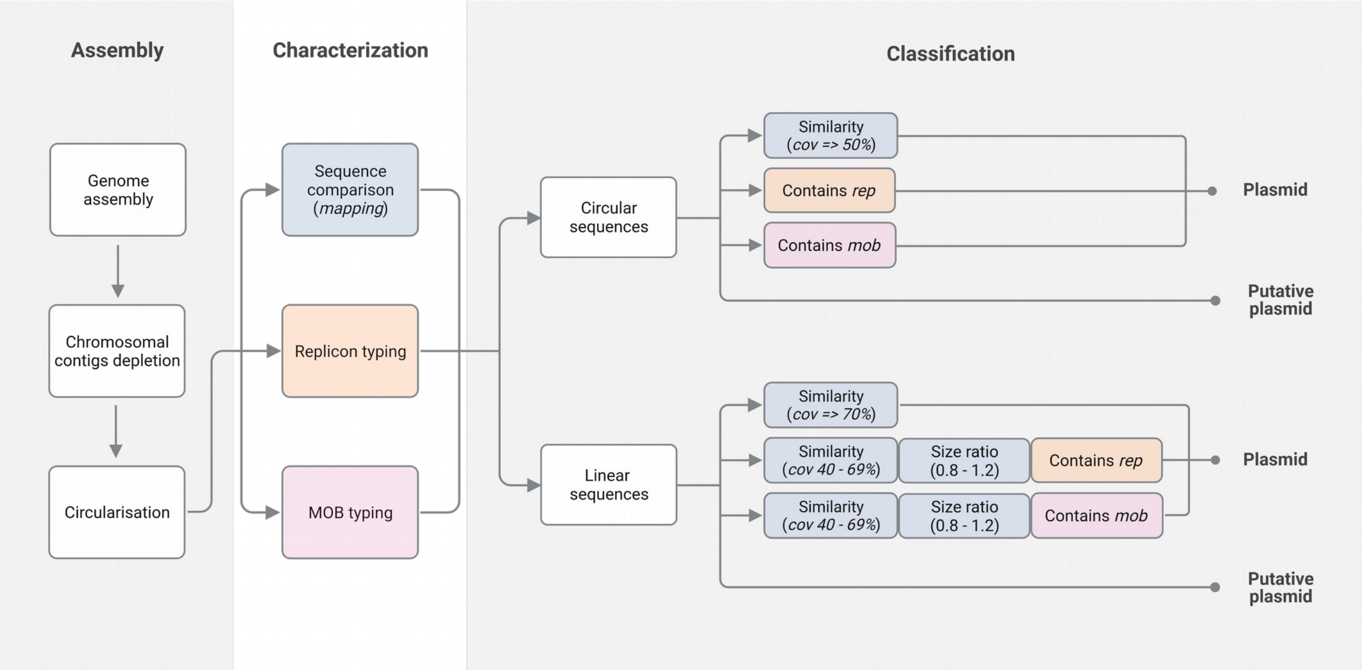


**Figure S13. Characterisation and classification of plasmid sequences from Murray genomes.** The figure illustrates the process followed for identifying plasmids from Murray genomes using a *mapping* strategy (See Methods for details). Contigs identified as plasmidic by plasmidverify ("Chromosomal contigs depletion" step) were further characterised as shown in the figure, including their comparison against known plasmids. Data gathered from the characterisation step was used to classify the sequences as plasmid or putative plasmid following the criteria indicated in the figure. Plasmid sequences detected with this strategy were combined with those identified by mob_recon to produce the final set of Murray plasmids reported in this study.

A

| Putative Murray plasmid sequence | Plasmid match | Plasmid match coverage |
|---|---|---|
| pMUR-M143_44 | CP055640.1 | 80% |
| pMUR-M614_30 | CP055640.1 | 100% |
| pMUR-M266_37.M266_38 | CP055640.1 | 100% |

**Escherichia coli strain RHB36-C13 plasmid unnamed**

GenBank: CP055640.1

FASTA Graphics

Go to:

```
LOCUS       CP055640                1941 bp    DNA     linear   BCT 16-MAY-2022
DEFINITION  Escherichia coli strain RHB36-C13 plasmid unnamed.
ACCESSION   CP055640
VERSION     CP055640.1
DBLINK      BioProject: PRJNA605147
            BioSample: SAMN15148367
KEYWORDS    .
SOURCE      Escherichia coli
  ORGANISM  Escherichia coli
            Bacteria; Pseudomonadota; Gammaproteobacteria; Enterobacterales;
            Enterobacteriaceae; Escherichia.
REFERENCE   1  (bases 1 to 1941)
  AUTHORS   AbuOun,M., Jones,H., Stubberfield,E., Gilson,D., Shaw,L.P.,
            Hubbard,A.T.M., Chau,K.K., Sebra,R., Peto,T.E.A., Crook,D.W.,
            Read,D.S., Gweon,H.S., Walker,A.S., Stoesser,N., Smith,R.P.,
            Anjum,M.F. and On Behalf Of The Rehab,Consortium.
  TITLE     A genomic epidemiological study shows that prevalence of
            antimicrobial resistance in Enterobacterales is associated with the
            livestock host, as well as antimicrobial usage
  JOURNAL   Microb Genom 7 (10) (2021)
   PUBMED   34609275
REFERENCE   2  (bases 1 to 1941)
  AUTHORS   Shaw,L.P.
  TITLE     Direct Submission
  JOURNAL   Submitted (12-JUN-2020) Nuffield Department of Medicine, University
            of Oxford, John Radcliffe Hospital, Oxford OX3 9DU, United Kingdom
COMMENT     The annotation was added by the NCBI Prokaryotic Genome Annotation
            Pipeline (PGAP). Information about PGAP can be found here:
            https://www.ncbi.nlm.nih.gov/genome/annotation_prok/

            ##Genome-Assembly-Data-START##
            Assembly Method       :: Unicycler v. v0.4.7
            Assembly Name         :: RH07 T3-C13
            Genome Representation :: Full
            Expected Final Version :: Yes
            Genome Coverage       :: 166.569649586864x
            Sequencing Technology :: Hybrid: Illumina and OXFORD NANOPORE
```

```
                     /country="United Kingdom"
                     /collection_date="2017"
     gene            <1..469
                     /locus_tag="HVZ50_23380"
     CDS             <1..469
                     /locus_tag="HVZ50_23380"
                     /inference="COORDINATES: similar to AA
                     sequence:RefSeq:NP_416064.1"
                     /note="Derived by automated computational analysis using
                     gene prediction method: Protein Homology."
                     /codon_start=2
                     /transl_table=11
                     /product="tail fiber assembly protein"
                     /protein_id="QLN25999.1"
                     /translation="GLPANSTDIAPPDIPAGFVAVFNSDKASWHLVEDHRGKTVYDVA
                     SGDALFISELGPLPENVTWLSPEGEFQKWNGTAWVKDAEAEKLFRIREAEETKNSLMQ
                     VASEHIAPLQDAVDLEIATEEETSLLEAWKKYRVLLNRVDTSTAQDIEWPALP"
     gene            complement(567..1157)
                     /locus_tag="HVZ50_23385"
     CDS             complement(567..1157)
                     /locus_tag="HVZ50_23385"
                     /inference="COORDINATES: similar to AA
                     sequence:RefSeq:NP_415892.1"
                     /note="Derived by automated computational analysis using
                     gene prediction method: Protein Homology."
                     /codon_start=1
                     /transl_table=11
                     /product="recombinase family protein"
                     /protein_id="QLN26000.1"
                     /translation="MSRIFAYCRISTLDQTIENQRREIESAGFKIKPQQIIEEHISGS
                     AATSERPGFNRLLARLKCGDQLIVTKLDRLGCNAMDIRKTVEQLTETGIRVHCLALGG
                     IDLTSPTGKMMMHVISAVAEFERDLLLERTHSGIVRARGAGKRFGRPPVLNEEQKQVV
                     FERIKSGVSISAIAREFKTSRQTILRAKAKLQTPDI"
     gene            complement(1474..1707)
                     /gene="ydfK"
                     /locus_tag="HVZ50_23390"
     CDS             complement(1474..1707)
                     /gene="ydfK"
                     /locus_tag="HVZ50_23390"
                     /inference="COORDINATES: similar to AA
                     sequence:RefSeq:NP_416062.2"
                     /note="Derived by automated computational analysis using
                     gene prediction method: Protein Homology."
                     /codon_start=1
                     /transl_table=11
                     /product="cold shock protein YdfK"
                     /protein_id="QLN26001.1"
                     /translation="MKSKDTLKWFPAQLPEVRIILGDAVVEVAKQGRPINTRTLLDYI
                     EGNIKKKSWLDNKELLQTAISVLKDNQNLNGKM"
     gene            complement(1776..1889)
                     /locus_tag="HVZ50_23395"
     CDS             complement(1776..1889)
                     /locus_tag="HVZ50_23395"
                     /inference="COORDINATES: similar to AA
                     sequence:RefSeq:YP_009518779.1"
                     /note="Derived by automated computational analysis using
                     gene prediction method: Protein Homology."
                     /codon_start=1
                     /transl_table=11
                     /product="hypothetical protein"
                     /protein_id="QLN26002.1"
                     /translation="MNSILIITSLLIIFSIFSHALIKLGIGISNNPDKTDV"
```

B

pMUR-M143_44

| | Description | Scientific Name | Max Score | Total Score | Query Cover | E value | Per. Ident | Acc. Len | Accession |
|---|---|---|---|---|---|---|---|---|---|
| ☑ | Escherichia coli strain RHB47-SO-C05 chromosome, complete genome | Escherichia coli | 66290 | 69966 | 97% | 0.0 | 99.96% | 4844648 | CP098929.1 |
| ☑ | Escherichia coli strain 18TN10VL01-EC chromosome, complete genome | Escherichia coli | 66290 | 70129 | 97% | 0.0 | 99.96% | 4690071 | CP063711.1 |
| ☑ | Escherichia coli strain RHB14-C21 chromosome, complete genome | Escherichia coli | 66290 | 69955 | 97% | 0.0 | 99.96% | 4898273 | CP057802.1 |
| ☑ | Escherichia coli strain RHB33-C12 chromosome, complete genome | Escherichia coli | 66290 | 69966 | 97% | 0.0 | 99.96% | 5026852 | CP057194.1 |
| ☑ | Escherichia coli strain RHB34-C14 chromosome, complete genome | Escherichia coli | 66290 | 70118 | 97% | 0.0 | 99.96% | 4644015 | CP057171.1 |
| ☑ | Escherichia coli strain NCTC9041 genome assembly, chromosome: 1 | Escherichia coli | 66284 | 70124 | 97% | 0.0 | 99.96% | 4693367 | LR134296.1 |

pMUR-M614_30

| | Description | Scientific Name | Max Score | Total Score | Query Cover | E value | Per. Ident | Acc. Len | Accession |
|---|---|---|---|---|---|---|---|---|---|
| ☑ | Escherichia coli strain RHBSTW-00673 chromosome, complete genome | Escherichia coli | 84195 | 94510 | 78% | 0.0 | 99.81% | 4787975 | CP056355.1 |
| ☑ | Escherichia coli strain RHB31-SO-C03 chromosome, complete genome | Escherichia coli | 83163 | 83357 | 74% | 0.0 | 99.41% | 4660104 | CP099123.1 |
| ☑ | Escherichia coli strain NCTC9066 genome assembly, chromosome: 1 | Escherichia coli | 82448 | 83945 | 74% | 0.0 | 99.13% | 4644134 | LR134000.1 |
| ☑ | Escherichia coli strain ETEC1723 chromosome, complete genome | Escherichia coli | 82010 | 1.040e+05 | 74% | 0.0 | 99.53% | 5708961 | CP122852.1 |
| ☑ | Escherichia coli strain ETEC6329F chromosome | Escherichia coli | 82003 | 99336 | 74% | 0.0 | 99.53% | 5218393 | CP122609.1 |

pMUR-M266_37.M266_38

| | Description | Scientific Name | Max Score | Total Score | Query Cover | E value | Per. Ident | Acc. Len | Accession |
|---|---|---|---|---|---|---|---|---|---|
| ☑ | Escherichia coli strain RHB34-SO-C08 chromosome, complete genome | Escherichia coli | 81061 | 1.696e+05 | 99% | 0.0 | 99.98% | 4739754 | CP099118.1 |
| ☑ | Escherichia coli strain FDAARGOS 1387 chromosome, complete genome | Escherichia coli | 81061 | 2.102e+05 | 100% | 0.0 | 99.98% | 5052411 | CP077379.1 |
| ☑ | Escherichia coli strain USDA-ARS-USMARC-51408 chromosome, complete genome | Escherichia coli | 81056 | 1.771e+05 | 99% | 0.0 | 99.98% | 4705478 | CP104114.1 |
| ☑ | Escherichia coli strain NCTC9100 genome assembly, chromosome: 1 | Escherichia coli | 81045 | 1.634e+05 | 96% | 0.0 | 99.98% | 4532589 | LR134239.1 |
| ☑ | Escherichia coli strain RHB41-E2-C01 chromosome, complete genome | Escherichia coli | 81045 | 1.850e+05 | 100% | 0.0 | 99.98% | 4751065 | CP099106.1 |
| ☑ | Escherichia coli strain 582 chromosome, complete genome | Escherichia coli | 81045 | 1.944e+05 | 95% | 0.0 | 99.98% | 4870922 | CP120568.1 |

**Figure S14. Examples of putative Murray plasmid sequences identified as contained in bacterial chromosomes.** Sequences from three Murray genomes originally classified as plasmids because of their similarity to plasmid CP055640.1 (**A**) were removed from further analysis because they also occur in chromosomal sequences (**B**).
